# TranSuite: a software suite for accurate translation and characterization of transcripts

**DOI:** 10.1101/2020.12.15.422989

**Authors:** Juan C. Entizne, Wenbin Guo, Cristiane P.G. Calixto, Mark Spensley, Nikoleta Tzioutziou, Runxuan Zhang, John W.S. Brown

**Affiliations:** Plant Sciences Division, School of Life Sciences, University of Dundee at The James Hutton Institute, Invergowrie, Dundee DD2 5DA, Scotland, UK; Information and Computational Sciences, James Hutton Institute, Dundee DD2 5DA, Scotland, UK; Donnelly Centre for Cellular and Biomolecular Research, University of Toronto, Toronto, Ontario, Canada; Cell and Molecular Sciences, James Hutton Institute, Dundee DD2 5DA, Scotland, UK

**Author notes:** Current address: Institute of Bioscience, University of São Paulo, São Paulo, Brazil (Cristiane P.G. Calixto). Co-corresponding authors Correspondence to: Dr. Runxuan Zhang, Prof John WS Brown.

**Keywords:** Coding sequence, open reading frame, translation start site, alternative splicing, premature termination codon, nonsense-mediated decay

## Abstract

Protein translation programs often select the longest open reading frame (ORF) in a transcript leading to numerous inaccurate and mis-annotated ORFs in databases. Unproductive transcript isoforms containing premature termination codons (PTCs) are potential substrates for nonsense-mediated decay (NMD). These transcripts often contain truncated ORFs but are incorrectly annotated due to selection of a long ORF beginning at an AUG downstream of the PTC despite the transcript containing the authentic translation start AUG. In gene expression and alternative splicing analyses, it is important to identify transcript isoforms which code for different protein variants and to distinguish these from potential NMD substrates. Here, we present TranSuite, a pipeline of bioinformatics tools that address these challenges by performing accurate translations, characterizing alternative ORFs and identifying NMD and other features of transcripts in newly assembled and existing transcriptomes. Directly comparing ORFs defined by TranSuite and TransDecoder for the Arabidopsis transcriptome AtRTD2 identified ORF mis-calling in over 16k (27%) of transcripts by TransDecoder.

## INTRODUCTION

In eukaryotes, translation initiation occurs by cap-dependent or cap-independent mechanisms. In cap-dependent initiation, a cap-binding complex recruits the mRNA and the 40S ribosomal subunit, which scans the 5’ UTR region for an AUG codon as the translation start site (Merchante et al., 2017). The AUG codon must have the proper context in the sequence to be accepted as the translation start site by the ribosome (Kozak, 1999, 2000; Lukaszewicz et al., 2000). The large ribosomal 60S subunit is recruited and translation proceeds. Upon encountering a stop codon, translation termination factors are recruited, the ribosomal subunits dissociate, thereby stopping translation and the mRNA and translated polypeptide are released. In contrast, cap-independent mechanisms, such as use of Internal Ribosome Entry Site (IRES), the ribosomal subunits directly engage with the translation start site within the mRNA. This mechanism is considered inefficient and rare for cellular mRNAs (Jackson, 2013). Similarly, in genes containing upstream Open Reading Frames (uORFs), re-initiation of translation at a downstream AUG or leaky scanning may occur but are inefficient and infrequent (Meijer and Thomas, 2002; Kochetov et al., 2008; Lee et al., 2012). In Arabidopsis, genome-wide ribosome profiling detected only 35 potential downstream alternative translation start sites in 31 genes (Liu et al., 2013). Ribosome profiling has also identified expressed uORFs which repress translation of downstream ORFs (Liu et al., 2013; Hsu et al., 2016). Therefore, although alternative translation mechanisms exist, in the majority of cases, the translation of transcripts in a gene will initiate with the same start codon and will terminate at the first stop codon.

Nonsense-mediated decay (NMD) is a translation-dependent mRNA surveillance pathway to detect and degrade aberrant mRNAs usually containing premature termination codons (PTC). Mechanisms and components of NMD in mammalian cells have been described in recent reviews (Hug et al., 2016; Karousis et al., 2016; Lykke-Andersen and Jensen, 2015; Nasif et al., 2018; Kurosaki et al., 2019). Aberrant RNAs can arise through mis-sense mutations and errors in transcription but a major source is alternative splicing (AS). NMD is important in regulating gene expression of physiologically relevant transcripts and AS isoforms and is modulated during development or in response to cellular or external environmental conditions (Karousis et al., 2016; Lykke-Andersen and Jensen, 2015; Hug et al., 2016; Nasif et al., 2018; Kurosaki et al., 2019). Many AS transcript isoforms contain PTCs and are potential targets of NMD (Schweingruber et al., 2013). Around 18% of the genes in Arabidopsis undergo AS resulting in unproductive mRNAs that contain PTCs and are degraded by NMD (Kalyna et al., 2012; Drechsel et al., 2013). PTCs are recognised by the presence of a downstream exon–exon junction located >50 nucleotides from the termination codon and deposition of an exon junction complex at the splice junction enhances NMD efficiency. Introns in 3’UTRs can induce NMD if >50 nt downstream of the authentic stop codon of mRNAs. PTCs lead to termination of translation by ribosomes at a distance from the polyA tail and such long 3’UTRs can activate NMD. The presence of short uORFs in 5’UTRs can also trigger NMD by initiating and terminating translation upstream of the translation start AUG of the main ORF. The rules governing the signals which target transcripts to NMD are not completely understood as not all PTCs, long 3’UTRs or uORFs trigger NMD (Karousis et al., 2016). Plants recognise PTC-containing transcripts by many of the same mechanisms as in mammals (Hori and Watanabe, 2007; Kertesz et al., 2006; Nyiko et al., 2009; Kalyna et al., 2012, Drachsel et al., 2013)). Plants also have differences in NMD activity in that PTC-containing transcripts due to retention of an intron are not susceptible to NMD (Kalyna et al., 2012; Marquez et al., 2012) because such transcripts appear to be retained in the nucleus and thereby avoid translation and NMD (Göhring et al., 2014). In addition, the presence of overlapping uORFs (a subclass of uORFs where the uORF overlaps the AUG of the main ORFs) correlated with NMD in Arabidopsis (Kalyna et al., 2012). AS in 5’ or 3’ UTRs can affect the size, position and nature of uORFs or the distance between the stop codon and splice junction of 3’UTR introns to impact NMD sensitivity of transcripts (Kalyna et al., 2012).

Accurate translation of mRNA sequences is an important aspect of analysing existing and newly assembled transcriptomes. Multiple programs are available for the identification, translation, annotation and characterization of ORFs, each with different functionalities. Widely used translation programs such as NCBI ORFfinder, EMBOSS getorf, and ExPASy Translate and more recent programs for the extraction of ORF sequences like OrfM (Woodcroft et al., 2016) translate all ORFs present in the transcripts. Other programs introduce methods to assess coding potential or other coding-related features. For example, CPC2 analyses ORF length and sequence composition (Kang et al., 2017), NMD Classifier measures the distance from a PTC/stop codon to the last splice-junction (Hsu et al., 2017), and sORFfinder identifies short ORFs with high coding potential (Hanada et al., 2010). Programs such as TransDecoder (https://github.com/TransDecoder/TransDecoder/wiki), the Bioconductor R packages ORFik (Tjeldnes and Labun, 2019) and SpliceR (Vitting-Seerup et al., 2014) allow identification of various features such as ORFs, uORFs, PTCs and NMD features and annotate CDS or ORF co-ordinates into a transcriptome annotation. The main disadvantages of these programs are that either that all ORFs in a transcript are reported requiring further analysis or they report the longest ORF which can be incorrect leading to mis-annotation. The programs vary greatly in ease of use, type and source of input data and the degree of downstream analysis required to extract meaningful information. Selection of the longest ORF in a transcript is accurate for many transcripts but is a simplistic approach given the complexity of features of mRNAs. In particular, thousands of alternatively spliced unproductive transcript isoforms that contain PTCs have mis-annotated ORFs in transcript annotations which are mis-leading for researchers (Brown et al., 2015). For example, when an AS event changes the reading frame of a transcript and introduces a PTC downstream of the authentic translation start site AUG, it is likely to leave a theoretical long ORF (>100 amino acids) further downstream in the transcript and it is this ORF that is identified by many translation programs. Thus, an ORF beginning at an AUG positioned downstream of the authentic translation start site is selected even though the transcript still contains the authentic translation start site AUG. As pointed out previously, a scanning ribosome has no ability to ignore the authentic translation start site in favour of a downstream AUG or ORF (Brown et al., 2015).

As more and more information on transcript diversity (alternative transcription start sites, alternative splicing and alternative polyadenylation) becomes available from short and long read RNA sequencing, it is necessary to accurately predict translations and characterize the protein-coding capacity of transcripts at the genome-wide level. To interpret gene expression and AS analyses, it is important to characterize transcripts and identify those which code for potentially functionally different protein variants and to distinguish those which are unlikely to code for proteins. Here, we present TranSuite, a program for identifying coding sequences, selecting translation start sites of a gene, accurately translating transcript isoforms and characterizing transcripts for their coding potential to identify protein variants and for features that likely trigger NMD. The advantages of TranSuite are that its approach to translation is more logical and is based on our knowledge of cellular mechanisms of translation, it brings together a range of protein and transcript analysis tools into a single, easy to use pipeline and provides a comprehensive information-rich output on gene and transcript characteristics.

## METHODS

### TranSuite pipeline

TranSuite consists of three independent modules: FindLORF, TransFix and TransFeat (Figure 1). FindLORF finds and annotates the longest ORF of each transcript; TransFix “fixes” the same translation start codon AUG in all the transcripts in a gene, and re-annotates the resulting ORF of the transcripts; TransFeat identifies structural features of transcripts, coding potential and NMD signals. The modules can be run independently or as a pipeline with a single command. Overall, all the ORFs in a transcript are identified and the longest ORF for each transcript is selected (FindLORF); all transcripts from the same gene are then compared and the translation start site AUG that generates the longest ORF is selected and used to translate all of the transcripts from that gene (TransFix); and finally, the translation information is used to define coding and non-coding genes and transcripts, coding potential, AS events in 5’ and 3’ UTRs, presence of PTCs and NMD features in transcript isoforms (TransFeat) (Figure 1).

**Figure 1.**
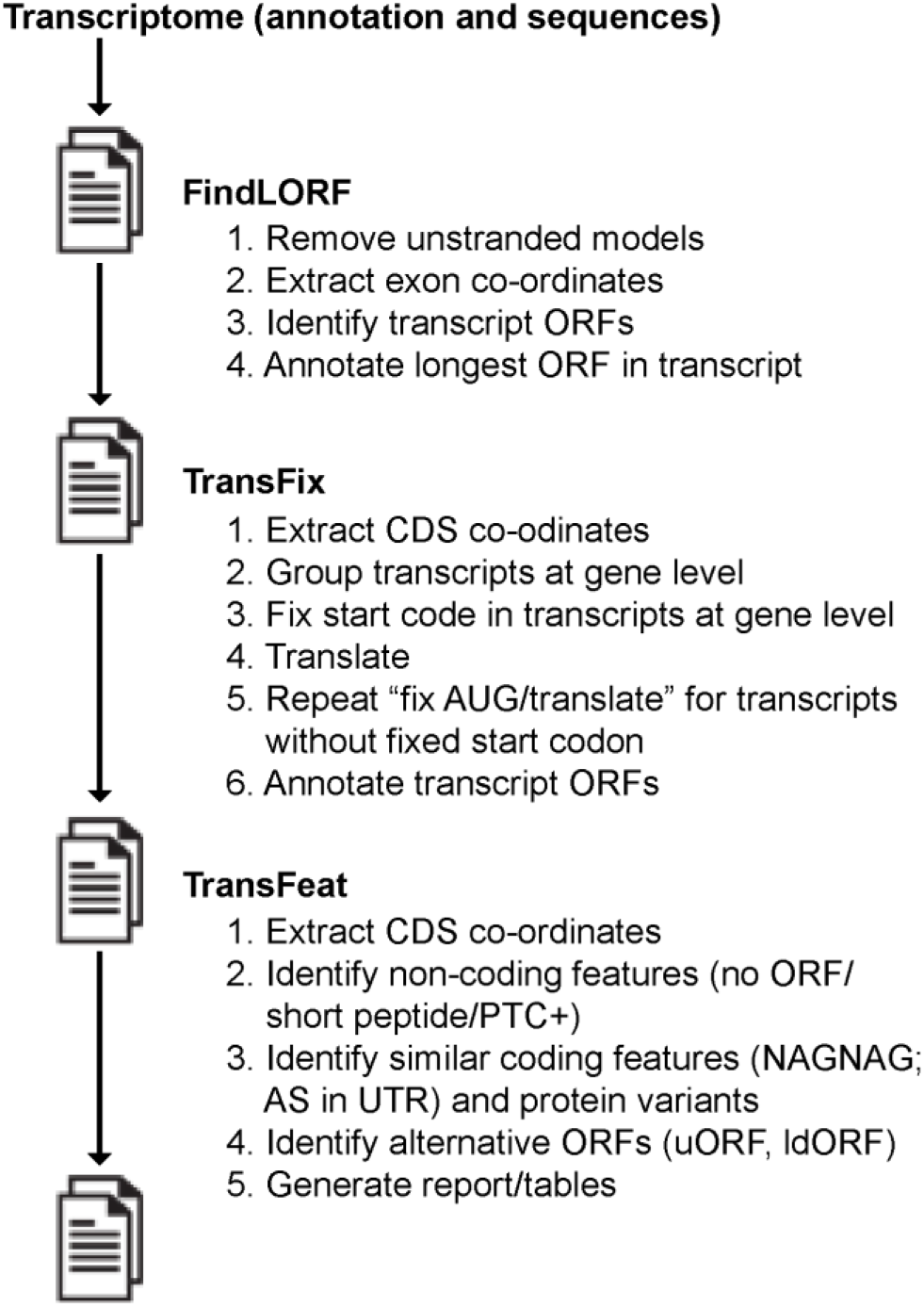
TranSuite pipeline. The TranSuite pipeline takes a) the transcripts to be analysed (e.g. from a new assembly or reference transcriptome) and b) their respective exonic sequences (FASTA file). *FindLORF* - For each transcript the longest ORF is identified and genomic co-ordinates generated. *TransFix* – all transcripts of a gene are translated with the “fixed” translation start AUG. *TransFeat* – transcript features are defined and transcripts classified. Information on genes and transcripts is contained in a TranSuite-generated report and tables.

### FindLORF

Transcriptome assemblers generate annotations containing only exon co-ordinate information. We implemented FindLORF to identify and annotate ORF/CDS information in newly assembled transcriptome annotations. Firstly, FindLORF translates each transcript sequence in its three frames of translation according to its annotated strand and stores the relative start and stop codon positions of all the resulting ORFs. Secondly, FindLORF selects the longest ORF for each transcript as its putative CDS region. Finally, FindLORF annotates the CDS using the genomic information contained in the transcriptome annotation to convert the relative ORF start-stop codon positions in the transcript sequence into genomic co-ordinates. The FindLORF module takes as input the transcriptome annotation to be curated (in GTF format) and a FASTA file containing the exon sequences of the transcripts. The output of FindLORF consists of the transcriptome annotation (GTF format) with the included CDS information, the FASTA files of the transcript sequences and their translations, as well as log files reporting transcripts that do not contain any start codons.

### TransFix

The TransFix module provides more biologically accurate translations by selecting the authentic translation start site for a gene, “fixing” this location and using it to translate the gene transcripts, and annotating the resulting CDS of the translations. We define the authentic translation start site as the site used to produce the full-length protein of the gene. In detail, TransFix firstly extracts the CDS co-ordinates of the transcripts from the transcriptome annotation and groups the transcripts according to their gene of origin. Then, TransFix selects the start codon of the longest annotated CDS in the gene as the representative (authentic) translation start site and translates all of the transcripts in the gene from the “fixed” translation start site. Finally, TransFix annotates the genomic co-ordinates of the resulting stop codons using the same approach described in FindLORF. In some cases, transcript isoforms do not contain the “fixed” translation start site due to an AS event or an alternative transcription start site (see Figure 4A and C). To account for this, TransFix tracks those transcripts that are not translated during the first fix AUG/translation cycle and they are then processed through a second fix AUG/translation cycle to determine and annotate their valid translation start-sites. Further rounds of fix AUG/translation are performed for a user-determined number of iterations (default=5). Notably, TransFix is also able to “fix” the translation start of chimeric genes. If the gene IDs of chimeric genes are provided, TransFix is able to select the “fixed” start codon of the 5’-most gene model to perform the fix AUG/translation cycle. The TransFix module takes as input the transcriptome annotation from FindLORF (above) and the FASTA file of the exon sequences of the transcripts. The output from TransFix is the transcriptome annotation with transcript and CDS structure information and the FASTA files of the transcript sequences and their translations.

### TransFeat

TransFeat extracts and processes the CDS information of transcripts contained in transcriptome annotations to infer multiple characteristics of the genes, transcripts and their coding potential (Figure 2), and reports the information in an easily accessible format. The key parameters used in TransFix are summarised in Table 1. TransFeat identifies non-coding and protein-coding transcript features. Non-coding transcripts either lack an annotated ORF (“No ORF”) due to the absence of an AUG or of any sufficiently long ORF (default: < 30 amino acids), or the annotated ORF is too short to be considered as coding (“Short ORF”) (default: < 100 amino acids). If a valid ORF is available, TransFeat proceeds to analyse those transcripts for coding-related features (Figure 2A). TransFeat then defines as “non-coding genes” those genes containing only non-coding transcripts (Figure 2A). All other genes are defined as protein-coding (Figure 2A). Genes with two or more transcript isoforms differing by at least one splice junction represent alternatively spliced genes. TransFeat characterises transcripts from protein-coding genes into isoforms that either code for protein variants or contain PTCs (Figure 2B). PTC-containing (PTC+) transcripts are defined as transcripts where the position of the ORF stop codon is below a threshold (default: 70%) relative to the longest ORF in the gene (which defines the 100% reference length). PTC+ transcripts from protein-coding genes represent unproductive transcripts because they do not code for full-length proteins and are likely NMD targets (Lareau et al., 2007). The transcripts that code for protein variants are then analysed to detect those that code for very similar proteins such as isoforms that differ by AS events occurring only in their UTR regions (“AS in UTR”) and isoforms whose translations differ by the addition/deletion of a single amino acid (“NAGNAG”) (Figure 2B). TransFeat detects the presence and location of AS in 5’ and/or 3’ UTR regions by grouping transcripts according to their exact CDS co-ordinates and comparing their mutual 5’ and 3’ UTR regions for differences of their intron co-ordinates. Transcripts with a single amino acid difference are identified by pairwise comparison of the CDS co-ordinates of the transcripts in a gene to detect those that differ by exactly 3 nt. The remaining transcript isoforms from protein-coding genes contain in frame AS events (exon skipping, intron retention, alternative 5’ and 3’ splice sites) which add or remove segments of coding sequence to give rise to protein variants (Figure 2B).

**Figure 2.**
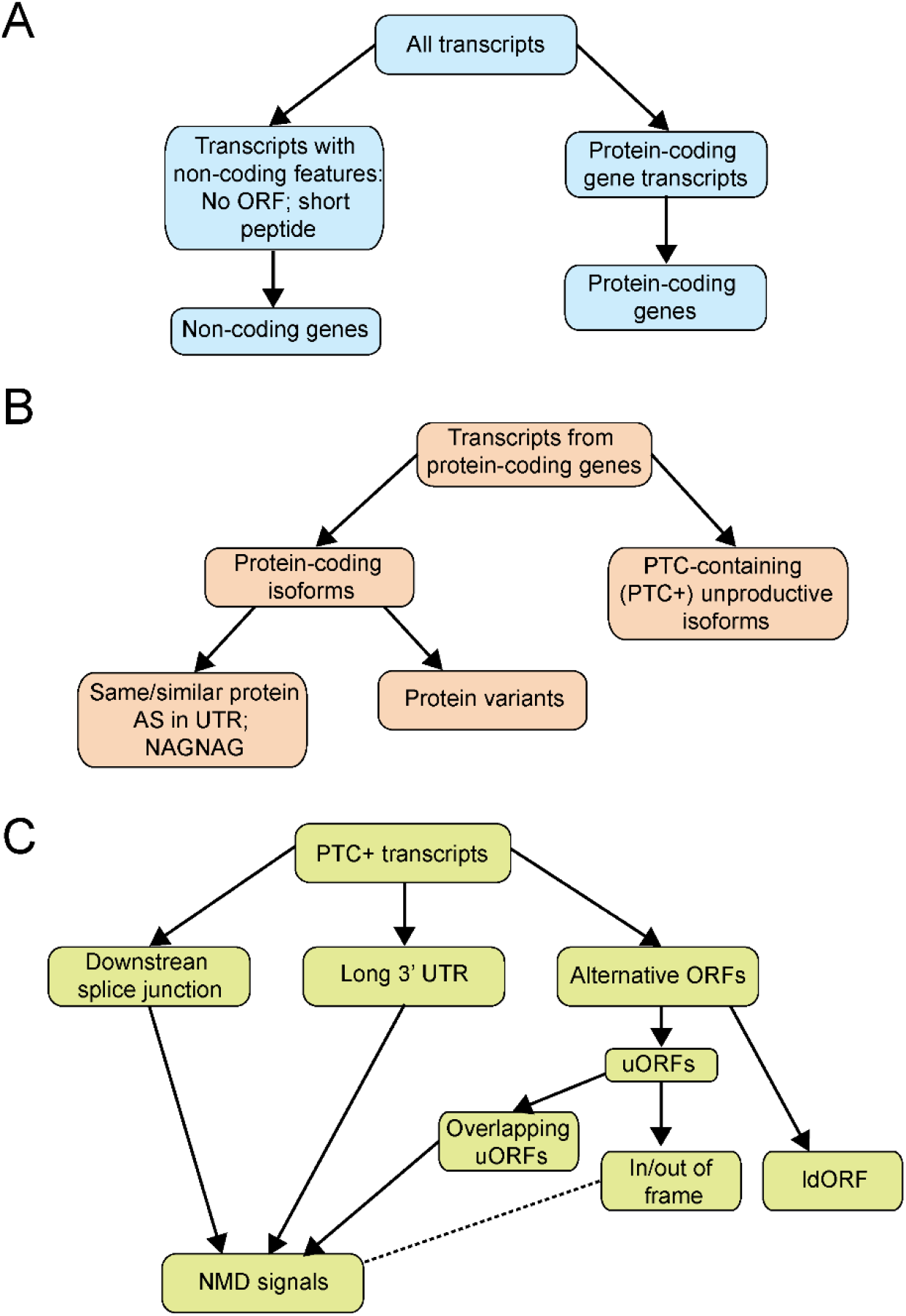
Transcript feature characterisation by TransFeat. A) translation data identifies transcripts with non-coding features (no ORF - no AUG; CDS < 30 amino acids or short peptide where CDS is between 30-100 amino acids (default values). Genes where all transcripts have non-coding features classed as Non-coding genes; all other genes (and their associated transcripts) are protein-coding genes. **B)** Transcripts containing a PTC (PTC+) are unproductive and all other transcripts are protein-coding isoforms. Transcripts of some genes code for identical proteins if AS occurs only in the 5’ and/or 3’UTRs. Transcripts with NAGNAG alternative splicing code for proteins which differ by one amino acid. Other protein-coding transcripts encode protein variants. **C)** PTC+ transcripts are further characterised by different features: downstream splice junction, long 3’UTR and overlapping uORF (NMD signals) as well as in frame and out of frame uORFs, long downstream ORFs. The dotted line from in/out of frame uORFs reflects the potential of some uORFs to trigger NMD.

**Table 1.** Transcript features identified by TransFeat and reported in the TranSuite output.

| **Feature** | **Description** |
| --- | --- |
| Coding | Protein-coding transcript isoform from a protein-coding gene |
| Non-coding | Transcript from a non-coding gene |
| Unproductive | Non-protein-coding transcript isoform from a protein-coding gene |
| No_ORF | Transcript does not contain an AUG or ORF is very short (default: <30 amino acids) |
| Short_ORF | Longest ORF less than threshold (default: <100 amino acids) |
| NAGNAG | NAGNAG or GTNGTN AS events - add or remove 3 nt and a single amino acid |
| AS_5UTR | AS in 5'UTR – does not affect CDS |
| AS_3UTR | AS in 3'UTR – does not affect CDS |
| NMD | Contains NMD features |
| PTC+ | Contains a premature termination codon |
| ds_SJ | Splice junction(s) >50nt downstream of stop codon/PTC |
| Long_3UTR | Long 3' UTR - distance from PTC to 3' end of transcript (default: >350 nt for Arabidopsis; Kalyna et al., 2012) |
| ouORF | uORF overlaps the translation start AUG |
| uORF | uORF upstream of the main ORF coding for a minimum of 10 amino acids |
| Inframe | uORF is in frame with the main ORF |
| Not_inframe | uORF is not in frame with the main ORF |
| ldORF | Long ORF downstream of a PTC |

Finally, TransFeat reports NMD-related features in PTC-containing transcripts such as the presence of splice junctions downstream of the stop codon (“dsSJ”), long 3’UTR regions (“long_3UTR”) and the presence of upstream ORFs overlapping the translation start site (““ouORF””) (Figure 2C) (Schweingruber et al., 2013; Kalyna et al., 2012). In addition, TransFeat detects and reports alternative ORF information using the same approach as detailed in FindLORF to translate all possible ORFs (default: minimum length 10 amino acids) and annotate their start-end co-ordinates. TransFeat uses this information to identify the presence of 1) upstream ORFs (“uORF”) in the 5’ UTR and whether they are in or out of frame with the main ORF of the gene for all protein-coding gene transcripts, and 2) long downstream ORFs (“ldORF”) for unproductive transcripts (Figure 2C). Upstream ORFs (uORFs) can trigger NMD but the features which determine whether a uORF impacts mRNA transcript stability are not well defined (Kalyna et al., 2012). PTC+ transcripts often contain a long ORF downstream of the PTC and have led to mis-annotation of many ORFs in databases (see Figure 3). TransFeat takes as input the transcriptome annotation with accurate CDS information, generated by TransFix, and the FASTA file of exon sequences of the transcripts. The output of TransFeat is a table with Gene/Transcript IDs, transcript features, CDS and genomic co-ordinates, and translations (Supplementary Table 1) and multiple files that allow the user to quickly explore the transcriptional landscape observed in the annotation (see Tables 3 and 4). Overall, the final outputs of the TranSuite pipeline are: 1) the corrected annotation generated by TransFix, 2) its respective transcript sequences and translation FASTA files, 3) TransFeat tables detailing the structural and coding information of transcripts and 4) multiple log files with detailed information of each step of the analysis. TranSuite summarises the numbers of coding and non-coding genes and transcripts, intron-containing and mono-exonic genes and genes with single and multiple transcript isoforms. It also reports more potential NMD signals than other available programs (overlapping uORF, long 3’ UTRs) and features of alternative ORFs (in or out of frame uORFs; ldORFs).

**Figure 3.**
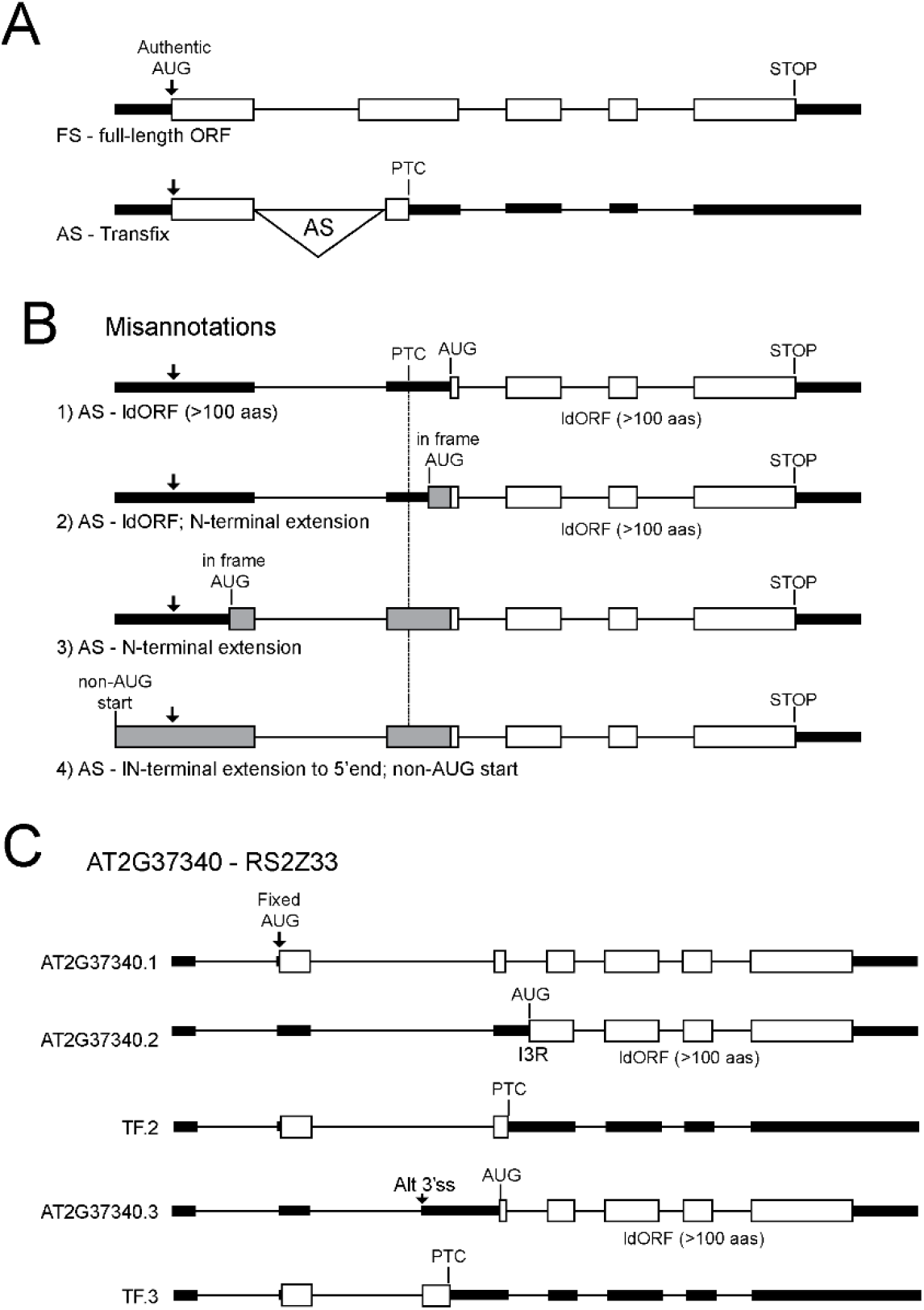
TransFix correctly translates and annotates ORFs in transcript variants. **A)** Schematic of gene with a fully spliced (FS) transcript that encodes the full-length protein and an alternatively spliced (AS) transcript where the AS changes the frame of translation and introduces a PTC (TransFix). **B)** Examples of mis-annotations of ORFs. 1) Mis-annotation of the AS transcript due to selection of the longest ORF defines an AUG downstream of the PTC (ignoring the authentic translation start site) producing a long downstream ORF (ldORF) which represents a C-terminal fragment of the protein of the gene. 2) Where an AUG in one of the other two reading frames occurs in frame and upstream of the ldORF, the ORF is extended (grey boxes). If this AUG is still downstream of the PTC, the ORF still represents an ldORF. 3) The AUG of the extended ORF can also be upstream of the PTC and 4) in TransDecoder, the ORF can extend to the 5’end of the transcripts. In 2), 3) and 4) the resultant ORFs are fusions of an N-terminal sequence (different from the protein coded by the gene) to a C-terminal region of the normal gene protein - all are incorrect. C) Transcripts of *RS2Z33* from TAIR10. The TAIR.2 and .3 transcripts are mis-annotated in TAIR and Araport as ldORFs but correctly annotated with TransFix as PTC-containing using the authentic (fixed) AUG which generates the RS2Z33 protein from the TAIR.1 transcript. White boxes – protein-coding exons; black boxes – UTRs; grey boxes – coding exons with sequence from different frame; arrow – position of authentic translation start AUG; in B) asterisk – PTC. AS events I3R in TAIR.2 and Alt 3’splice site (Alt 3’ss) in TAIR.3 generate PTCs (shown in the TF transcripts) but are mis-annotated to contain downstream ORFs in TAIR.2 and .3.

### AtRTD2 analysis with TranSuite and TransDecoder

The Arabidopsis AtRTD2 transcriptome (Zhang et al., 2017) was analysed with TranSuite and TransDecoder to identify and annotate the CDS of its transcripts and translate them. TranSuite analysis was run using its Auto module which automatically runs the whole pipeline (Figure 1). The analysis was run using the default parameters (see Table 1) with the exception of specifying a minimum of 70 AA to consider an ORF as potentially “coding” (see “Short_ORF”, Table 1). TransDecoder analysis was run as per the guidelines on the TransDecoder website: “Starting from a genome-based transcript structure GTF file”. The analysis was run with the default parameters, with the exception of specifying to retain only the single best ORF per transcript (--single_best_only) during the TransDecoder.Predict step. The code of the analysis is provided in our GitHub repository (https://github.com/anonconda). Transcripts with translations were viewed in the Integrative Genomics Viewer (IGV) from the Broad Institute.

## RESULTS

### TranFix overcomes mis-annotation of ORFs in transcript models

Many transcript models in databases are mis-annotated in terms of their coding potential because the longest ORFs have been selected irrespective of position in a transcript (Brown et al., 2015). When AS events shift the reading frame to introduce a PTC, its position will affect other ORFs in the transcript. For example, a transcript with a PTC towards the 5’ end of the CDS is more likely to contain a long ORF downstream of the PTC (long downstream ORF; “ldORF”), despite the transcript containing the authentic translation start site AUG that would normally be used to translate mRNAs coding for the full-length protein (Figure 3A). In contrast, a PTC towards the 3’-end of a transcript is likely to change the reading frame giving a protein with a different C-terminal region. The advantage of TransFix is that it uses the same AUG to translate all transcripts from the same genomic locus, thereby correctly identifying protein-coding transcript isoforms and distinguishing them from unproductive transcripts which contain PTCs. In contrast, translation programs that identify the longest ORF ignore the presence of authentic translation start sites and often predict long downstream ORFs (Figure 3B). The nature of such predicted ORFs also varies dependent on the position of AUGs in other reading frames. For example, downstream of the PTC, the longest ORF is likely to be a C-terminal fragment of the normal protein sequence of the gene (Figure 3B.1). However, if an AUG in one of the other two reading frames is now in frame with the ldORF, the predicted ORF will consist of the C-terminal fragment fused to a peptide sequence at the N-terminal end that is unrelated to the normal protein sequence of the gene as it comes from one of the other reading frames (Figure 3B.2 and .3).

As an example, the mis-annotation of ldORFs is illustrated for AtRS2Z33 (AT2G37340), an Arabidopsis SR splicing factor that has six transcript isoforms, only one of which codes for the full-length RS2Z33 protein (Figure 3C). The .2 and .3 isoforms illustrate mis-annotation of ORFs in TAIR10 and Araport11. The AT2G37340.2 transcript has retention on intron 3 (I3R) and the AT2G37340.3 transcript has an alternative 3’ splice site in intron 2 which leaves 218 nt of the intron sequence in the transcript. The annotated ORF in the .2 transcript in TAIR and Araport uses an AUG within the retained intron sequence and that in the .3 transcript uses a different AUG in exon 3; both transcripts generate ORFs consisting of C-terminal fragments of the CDS but still contain the translation start site sequence used to translate the protein from the AT2G37340.1 transcript. Using Transfix, the AUG in AT2G37340.1 (longest ORF) was fixed and all three transcripts were translated from this AUG. The .2 transcript ORF now stops at a stop codon in the retained intron (TF.2) while in the .3 transcript, the ORF stops within the part of intron 2 which remains after the AS event (TF.3). Both transcripts are therefore unproductive. They contain PTCs and have multiple downstream splice junctions. As pointed out previously (Brown et al., 2015) there is no known mechanism for ribosomes to ignore the normally used AUG and preferentially select a downstream AUG because it could lead to a longer downstream ORF. Indeed, one of the features identified by TransFeat are transcripts which contain such long downstream ORFs to help to identify such mis-annotations and avoid mis-interpretation.

### Translation of transcripts missing the first fixed AUG

Some protein-coding genes have transcripts where the fixed start codon is absent, and an alternative AUG is used to generate an N-terminally truncated protein. For example, genes with alternative transcription start sites (Figure 4A) and genes with AS events that remove the authentic AUG in some transcripts (Figure 4C) generate transcripts without the fixed AUG. To address these situations, TransFix performs multiple iterative “fix AUG/translation” cycles. That is, transcripts that are not translated in the first round are recycled to undergo another round of identifying the longest ORF, fixing the AUG and translating the remaining transcripts. In the translation of Arabidopsis AtRTD2, over 94% of transcripts were translated in the first cycle and 5% in the second cycle of translation (Table 2). At each cycle, the module tracks whether a transcript has been successfully translated and the remaining transcripts are processed in a new cycle. For example, AT1G18390 (*LRK10L1 – LEAF RUST RECEPTOR-LIKE KINASE 10-LIKE 1*) has two transcripts produced from alternative transcription start sites (Shin et al., 2015). TransFix identifies and fixes the AUG for the AT1G18390.1 transcript (Figure 4B). Transcription of AT1G18390.2 starts in intron 1 relative to the gene model; this transcript lacks the fixed AUG in AT1G18390.1 and enters a second round of translation which identifies a second AUG for translation (Figure 4B). AT5G09230 (*SIRTUIN 2 HISTONE DEACETYLASE family* - *SRT2*) contains numerous transcripts. Comparing AT5G09230.5 to AT5G09230.1, an AS event in .5 skips exon 2 which contains the AUG fixed in the first cycle of translation (Figure 4D). AT5G09230.5 enters a second cycle of translation and a second AUG which generates the longest ORF for this transcript is identified (Figure 4D). The AT5G09230.4 is a further example of a mis-annotated ORF. It contains an AS event that introduces a PTC but a long downstream ORF is mis-annotated in TAIR10 and Araport11. TransFix uses the fixed AUG from AT5G09230.1, generating a truncated ORF (TF.4) (Figure 4D).

**Figure 4.**
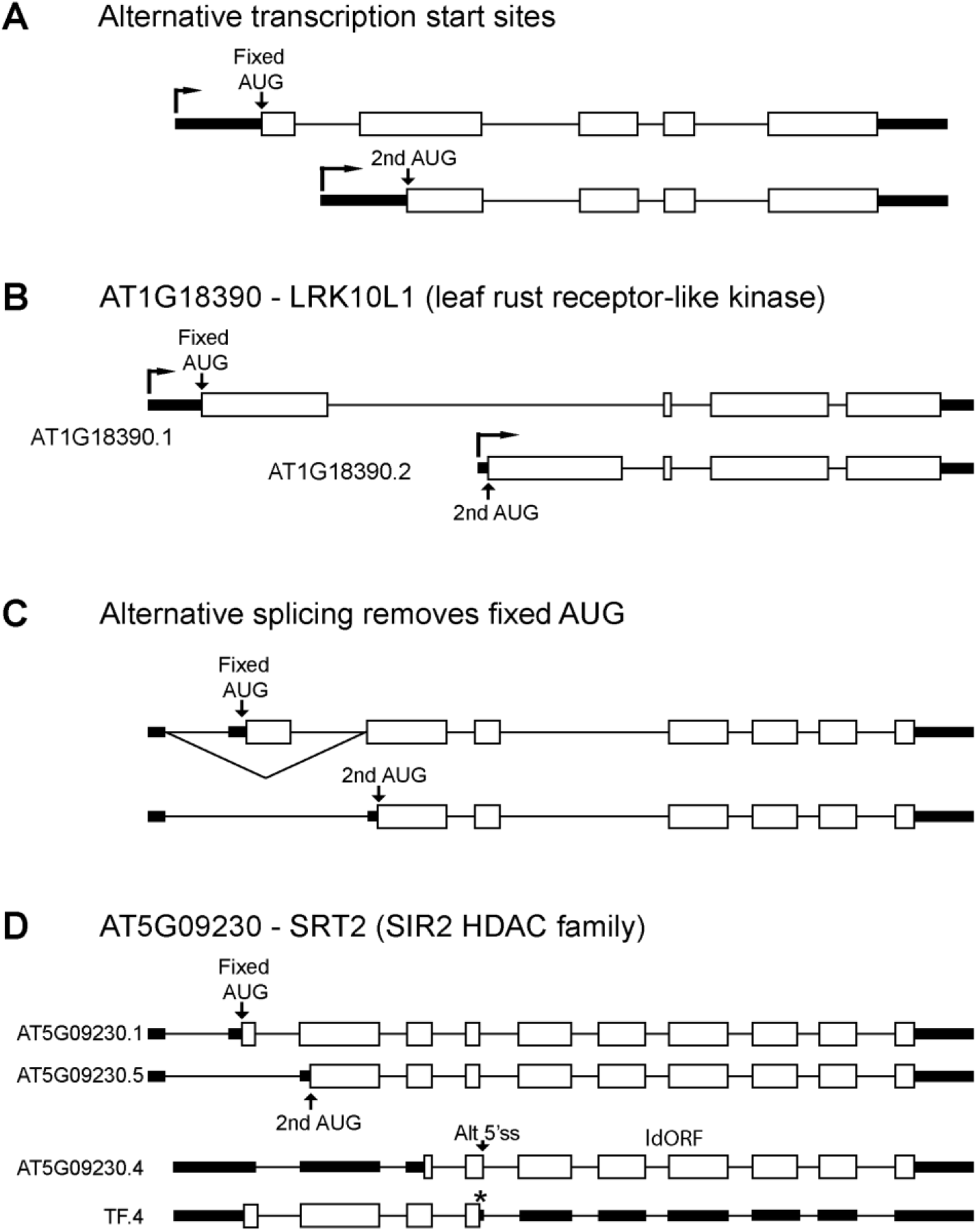
Translation of transcripts with alternative transcription start sites or where AS removes fixed AUG. **A)** Alternative transcription start sites: the fixed AUG of the longer transcript is absent in the shorter transcript due to a different TSS; the shorter transcript is recycled through TransFix a second time to identify the second AUG which produces the longest ORF for this transcript. **B)** Example of gene with different TSS. The fixed AUG in AT1G19390.1 is not present in AT1G13930.2 which begins within intron 1. TransFix identifies the second AUG used for translation of this transcript. **C)** Alternative splicing removes the fixed AUG. TransFix fixes the AUG in exon 2 of the upper transcript that codes for the full-length protein; the lower transcript has an AS event (exon skipping) that removes exon 2 and the fixed AUG. The lower transcript enters a second cycle of TransFix and a second AUG is fixed and used for translation. **D)** Example of AS removing a fixed AUG. TransFix fixes the AUG in exon 2 of AT5G09230.1 which is absent in AT5G09230.5 due to an exon skipping event; a second cycle of TransFix identifies the AUG for translation of this transcript isoform. AT5G09230.4 is a further example of a transcript with mis-annotated ORF in TAIR10 and Araport11 corrected by TransFix (TF.4). White boxes – protein-coding exons; black boxes – UTRs; angled arrows – transcription start site; fixed and second AUGs are indicated; asterisk – PTC; Alt 5’ss – alternative 5’ splice site.

**Table 2.** Number of transcripts of AtRTD2 translated in each consecutive cycle of translation.

| **Cycle number** | **Number of transcripts** | **% of total transcripts** |
| --- | --- | --- |
| 1 | 77137 | 93.9 |
| 2 | 4334 | 5.3 |
| 3 | 304 | 4.0 |
| 4 | 22 | 0 |
| 5 | 2 | 0 |

### TranSuite analysis of Arabidopsis AtRTD2 transcriptome

The Arabidopsis AtRTD2 transcriptome (Zhang et al., 2017) contains 82,190 unique transcripts (34,212 genes). In the first cycle of translation ca. 94% of transcripts were successfully translated or were identified as No ORF or Short ORF transcripts (Table 2). The second cycle of translation translates the majority of the remaining transcripts (4,334; 5.3% of total) which represented transcripts from genes with alternative transcription start sites or where an AS event removed the first fixed AUG (Figure 4A and C). The output from TranSuite consists of summary tables of genes and transcripts classified as single or multi-exonic genes with single or multiple transcript isoforms (Table 3) and by transcript coding and NMD features (Table 4). For AtRTD2, 84.5% of genes were protein-coding and 15.5% were non-coding (Table 4; Figure 5A); 70% were multi-exonic and 30% were mono-exonic (Table 3; Figure 5B). In terms of the number of transcript isoforms per gene, 44% of all genes were multi-exonic with multiple isoforms (reflecting AS); 26% and 30% were multi-exonic or mono-exonic genes producing single transcripts, respectively (Table 3; Figure 5C). For protein-coding genes only, half were multi-exonic with multiple isoforms and half had a single transcript from both multi- and mono-exonic genes (Table 4; Figure 5D). Of the 5,314 non-protein-coding genes, 558 were multi-exonic with multiple isoforms and therefore undergo AS.

**Figure 5.**
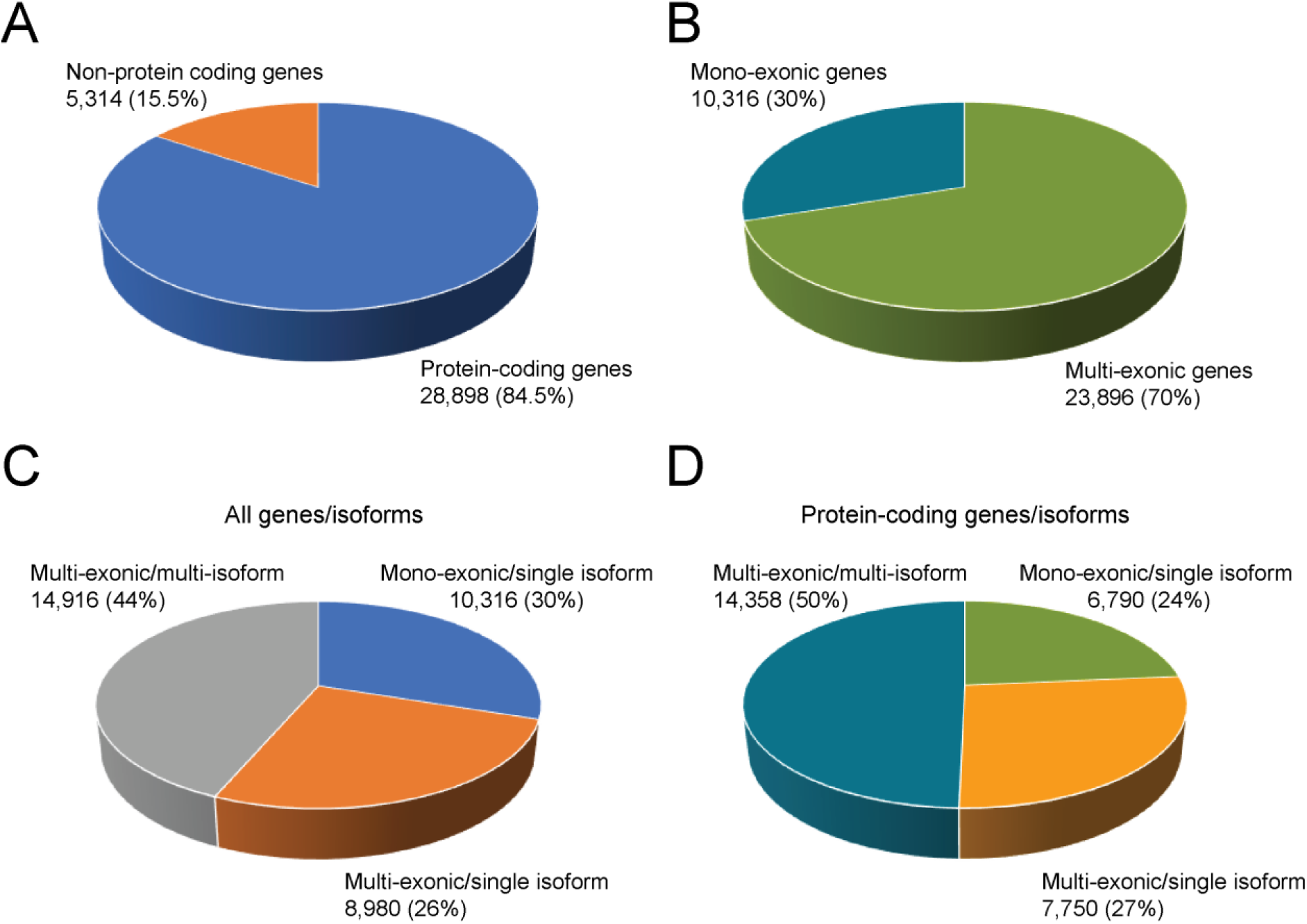
TranSuite analysis of gene classes and distribution of isoforms in Arabidopsis AtRTD2 transcriptome. The 34,212 genes of AtRTD2 are divided into **A)** protein-coding and non-protein-coding genes; and **B)** mono-exonic and multi-exonic genes. Distribution of isoform numbers in mono-exonic and multi-exonic genes for **C)** all genes, and **D)** protein-coding genes. The analysis is based on gene classification using a minimum of 100 amino acids to define a protein-coding transcript.

At the transcript level, 54,425 of the 82,190 (66.2%) transcripts in AtRTD2 were protein-coding, 25.7% were unproductive transcripts from protein-coding genes and 8.1% were from non-protein-coding genes (Table 4; Figure 6A). Protein-coding transcripts were divided into transcripts which coded for identical proteins or proteins differing by a single amino acid. Transcript isoforms coding for the same protein have AS events only in UTRs (AS UTR) and those by only 1 amino acid had NAGNAG (or GTNGTN) AS events where alternative splicing removed only 3 nt (AS UTR and NAGNAG made up 29.6% of transcripts). NAGNAG events are relatively common in plants (Schindler et al., 2008; Sinha et al., 2010). The remaining transcripts (70.4%) encoded protein variants (Table 4; Figure 6B). The NAGNAG/AS UTR transcripts were further broken down into whether AS events were in the 5’ and/or 3’UTR or were NAGNAG/GTNGTN (Table 3; Figure 6C). The most frequent AS events were in the 5’UTR (66%) occurring around three times more frequently than in the 3’UTR. NAGNAG/GTNGTN AS events were present in 3.4% of all transcripts. Finally, TransFeat provides comprehensive information of the range of different NMD features of the unproductive transcripts from protein-coding genes (Table 4). In particular, the vast majority (ca. 76%) of the unproductive transcripts contained classical NMD features of a PTC with splice junctions >50 nt downstream and long 3’UTRs and a further ca. 16% of transcripts also having an overlapping uORF or either a DsSJ or long 3’UTR (Table 4; Figure 6D). Thus, TranSuite generates a detailed set of information on the genes and transcripts in an annotation both in terms of protein coding capacity and NMD features based on accurate translations.

**Table 3.** Summary of TranSuite output of AtRTD2 for mono-exonic/multi-exonic genes with single or multiple transcript isoforms.

|  | Number<br>(70 AA)* | Percentage<br>(70 AA)† | Number<br>(100 AA)* | Percentage<br>(100 AA)† |
| --- | --- | --- | --- | --- |
| **SINGLEISO_MONOEXONIC** |  |  |  |  |
| **GROUP_TOTAL** |  |  |  |  |
| GROUP_GENES | 10316 | 30.2 | 10316 | 30.2 |
| GROUP_TRANS | 10316 | 12.6 | 10316 | 12.6 |
| **GENE_SUBCATEGORIES** |  |  |  |  |
| CODING_GENES | 7799 | 22.8 | 6790 | 19.8 |
| NONCODING_GENES | 2517 | 7.4 | 3526 | 10.3 |
| **TRANS_SUBCATEGORIES** |  |  |  |  |
| CODING_TRANS | 7799 | 9.5 | 6790 | 8.3 |
| NONCODING_TRANS | 2517 | 3.1 | 3526 | 4.3 |
| **SINGLEISO_MULTIEXONIC** |  |  |  |  |
| **GROUP_TOTAL** |  |  |  |  |
| GROUP_GENES | 8980 | 26.2 | 8980 | 26.2 |
| GROUP_TRANS | 8980 | 10.9 | 8980 | 10.9 |
| **GENE_SUBCATEGORIES** |  |  |  |  |
| CODING_GENES | 8475 | 24.8 | 7750 | 22.7 |
| NONCODING_GENES | 505 | 1.5 | 1230 | 3.6 |
| **TRANS_SUBCATEGORIES** |  |  |  |  |
| CODING_TRANS | 8475 | 10.3 | 7750 | 9.4 |
| NONCODING_TRANS | 505 | 0.6 | 1230 | 1.5 |
| **MULTIISO_MULTIEXONIC** |  |  |  |  |
| **GROUP_TOTAL** |  |  |  |  |
| GROUP_GENES | 14916 | 43.6 | 14916 | 43.6 |
| GROUP_TRANS | 62894 | 76.5 | 62894 | 76.5 |
| **GENE_SUBCATEGORIES** |  |  |  |  |
| CODING_GENES | 14685 | 42.9 | 14358 | 42 |
| NONCODING_GENES | 231 | 0.7 | 558 | 1.6 |
| **TRANS_SUBCATEGORIES** |  |  |  |  |
| CODING_TRANS | 40937 | 49.8 | 39885 | 48.5 |
| UNPROD_TRANS | 21234 | 25.8 | 21132 | 25.7 |
| NONCODING_TRANS | 723 | 0.9 | 1877 | 2.3 |

\*70AA and 100AA refer to upper limit defining a short peptide; † Percentage of total genes or transcripts

**Table 4.**
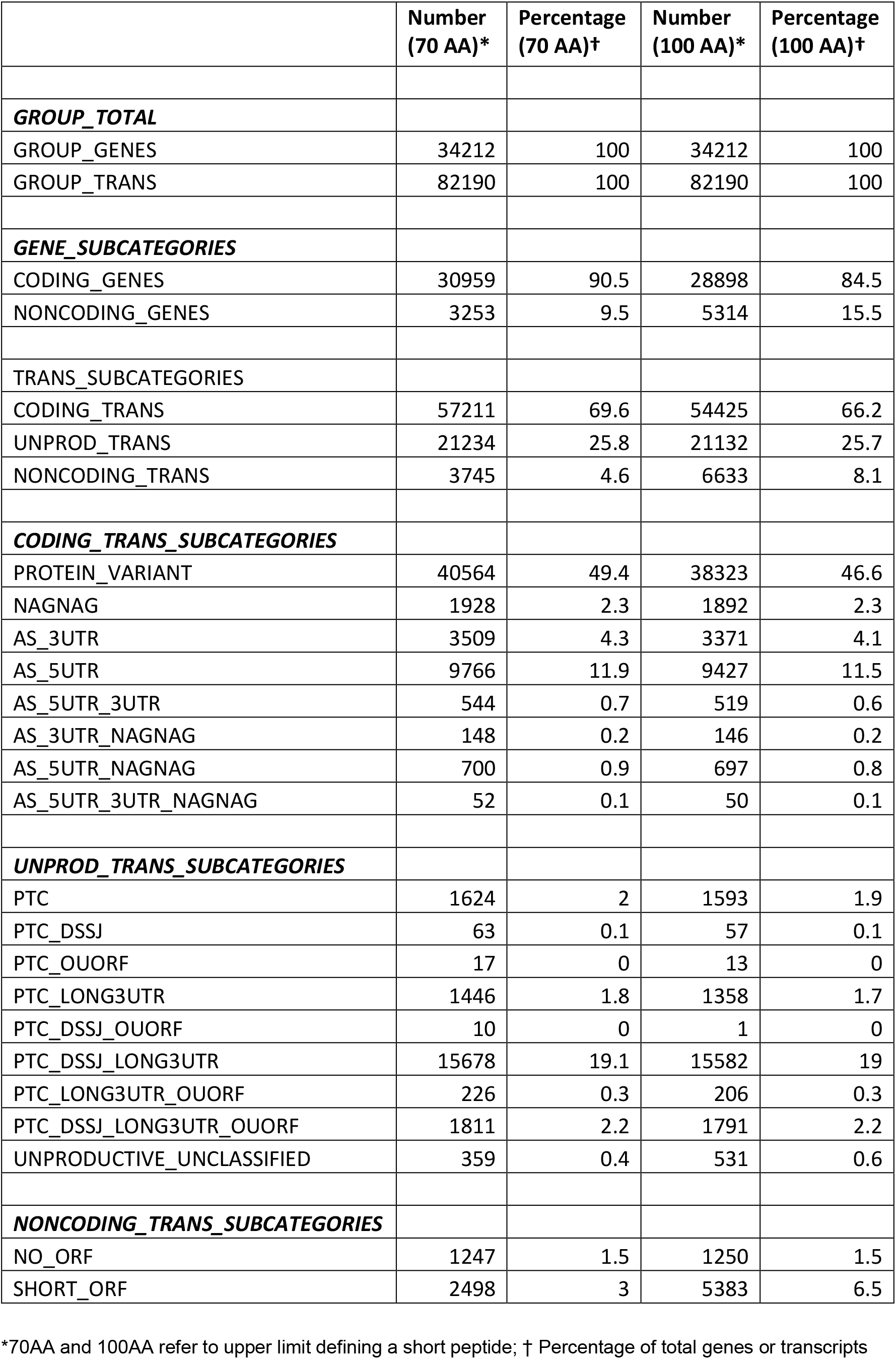
Summary of TranSuite analysis on AtRTD2 gene/transcript characterisation.

**Figure 6.**
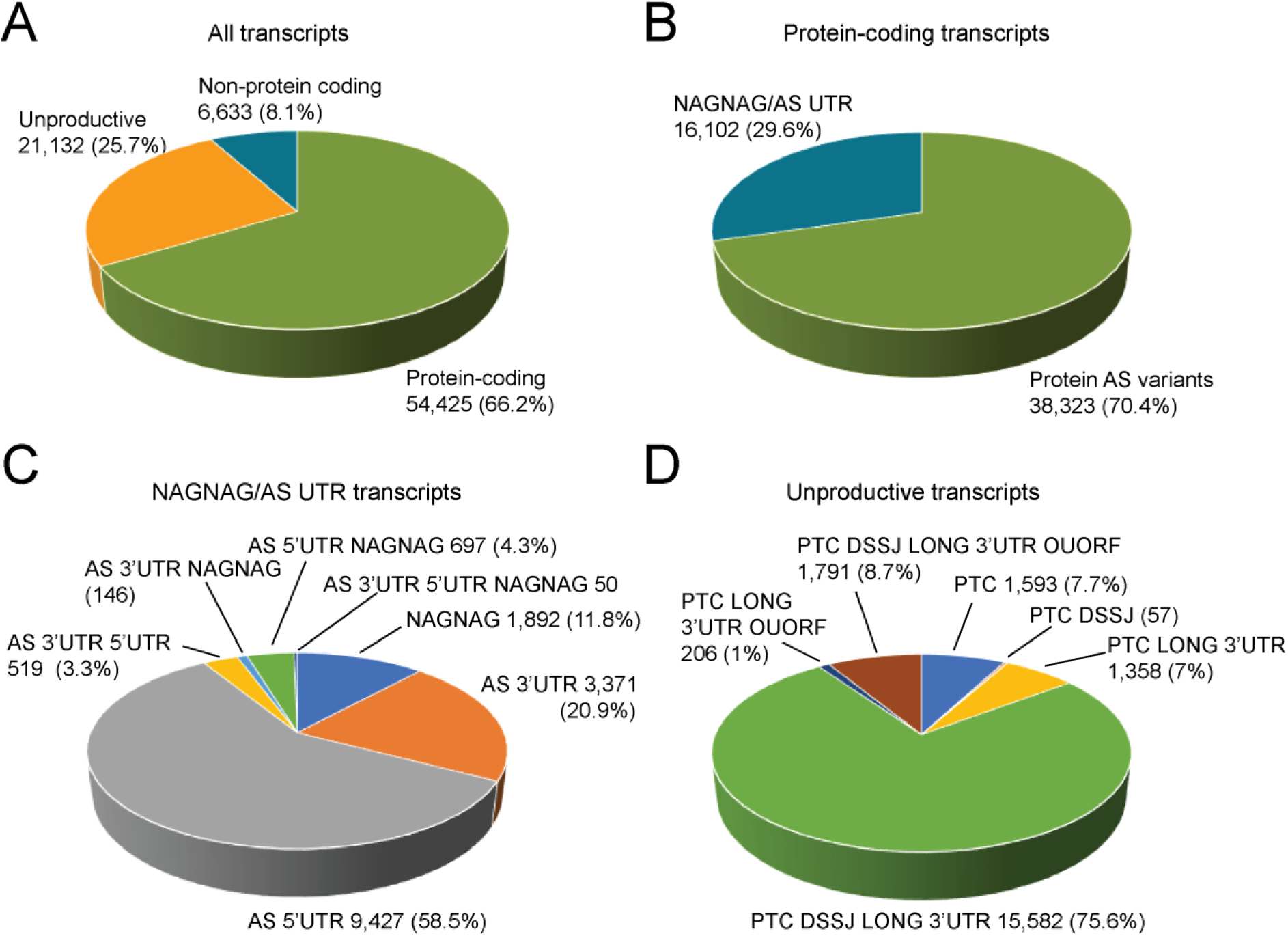
TranSuite analysis of Arabidopsis AtRTD2 transcriptome. **A)** The 82,190 transcripts of AtRTD2 are divided into transcripts from protein-coding genes (both protein-coding and unproductive transcripts) and those from non-protein-coding genes; **B)** protein-coding transcripts are separated into those transcripts with no or little change in protein sequence due to AS occurring only in the 5’ and/or 3’UTR or NAGNAG AS which alters the coding sequence by a single amino acid, and those coding for protein variants; **C)** NAGNAG and AS UTR transcripts are divided into different combinations of these AS events; **D)** unproductive transcripts which contain PTCs are divided by the presence of different NMD features (DSSJ – downstream splice junction; Long 3’UTR; OUORF – overlapping uORF. The analysis is based on gene classification using a minimum of 100 amino acids to define a protein-coding transcript.

### Comparison of translations by TranSuite and TransDecoder of Arabidopsis AtRTD2 transcriptome

Multiple programs are available for the identification, translation, annotation and characterization of transcripts ORFs. The different functionalities and features of a number of such programs were compared to TranSuite to illustrate differences among programs and the more comprehensive nature of TranSuite (Table 5). Currently, TransDecoder is one of the most widely used programs for the identification, translation and annotation of CDS regions. It is similar in scope, functionality and ease of use to TranSuite, and importantly, like TranSuite, it generates a transcriptome annotation with the predicted ORFs annotated using genomic co-ordinates. However, the major difference is that TransDecoder identifies ORFs longer than a minimum length (e.g. >100 amino acids) and applies a scoring system which prioritises ORFs in the first reading frame (same frame as identified ORF) and by the length of the ORF. In contrast, TranSuite translates transcripts using a fixed translation start site which represents the authentic start site. The consequence is that TransDecoder is more likely to select ldORFs while TranSuite will report truncated ORFs ending in a PTC. To compare ORF prediction by TranSuite and TransDecoder directly, we provided the same input to both analyses: 1) the AtRTD2 transcriptome annotation (Zhang et al., 2017) without CDS information and 2) the Arabidopsis Genome FASTA file, to generate the transcriptome FASTA file of AtRTD2. We compared the CDS co-ordinates between the output from TransFix and TransDecoder to identify exact matches (Figure 7A). Over half of the ORFs (ca. 44k - 54%) had identical co-ordinates and represented ORFs from transcripts of mono- and multi-exonic protein-coding genes with a single isoform, transcripts from multi-isoform genes which code for full-length protein variants and transcripts from genes where AS occurs only in the 5’ or 3’ UTR. The main differences were that ca. 21k (25.60%) and 16k (20.04%) of transcripts were unique to TransFix and TransDecoder, respectively (Figure 7A). We expect that ORFs unique to TranSuite will largely reflect truncated ORFs ending in a PTC and is consistent with AtRTD2 containing ca. 21k PTC+ unproductive transcripts from protein-coding genes (Table 4). The majority of these will generate truncated ORFs in TransFix while many will produce ldORFs in TransDecoder. Therefore, the prediction from the different methods of ORF determination is that TransDecoder should report a higher number of longer ORFs than TransFix. Plotting ORF length derived from each transcript clearly shows the higher number of longer ORFs produced by TransDecoder (Figure 7B).

**Table 5.**
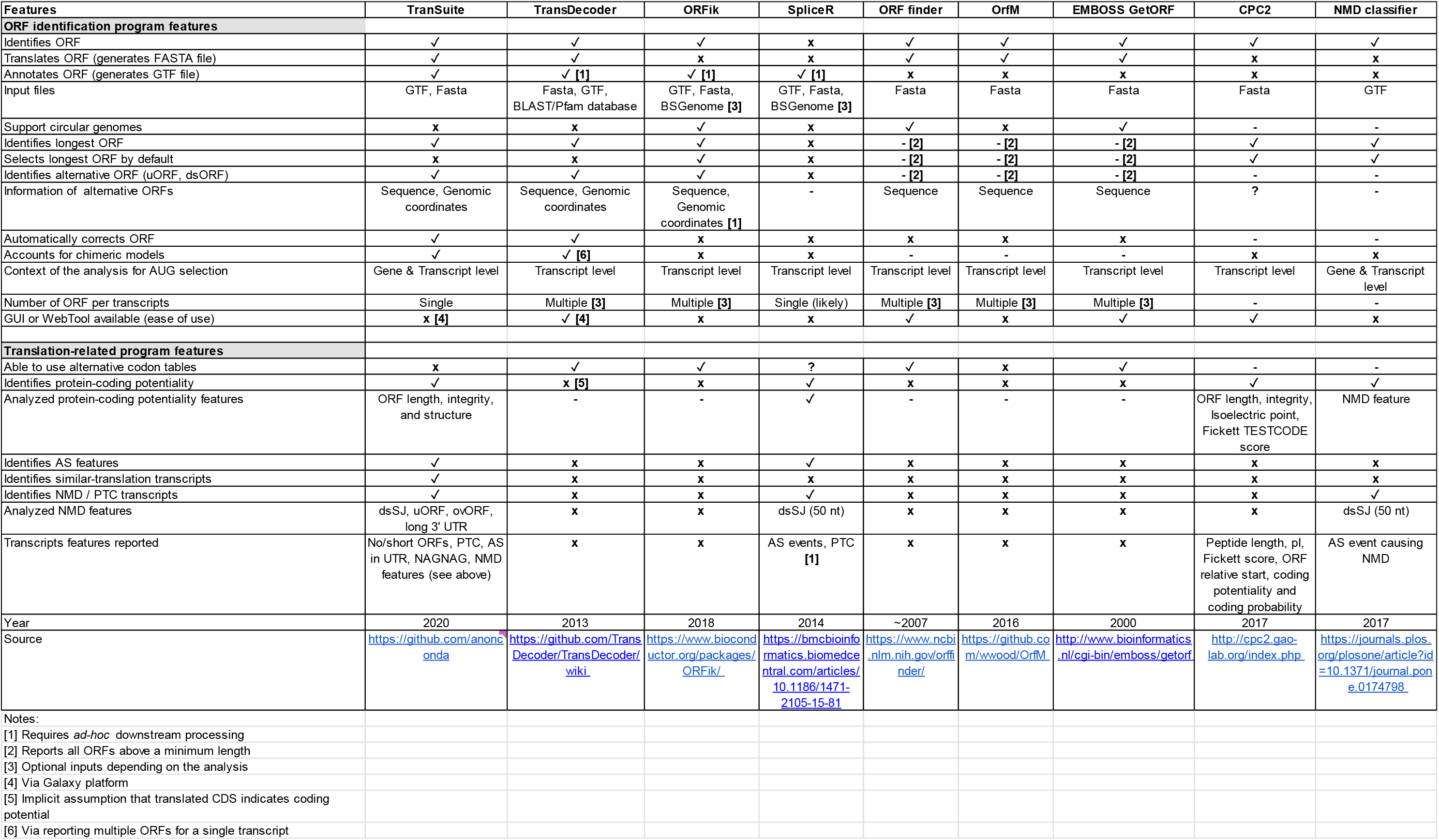
Comparison of features of TranSuite to other translation and AS/NMD feature programs.

**Figure 7.**
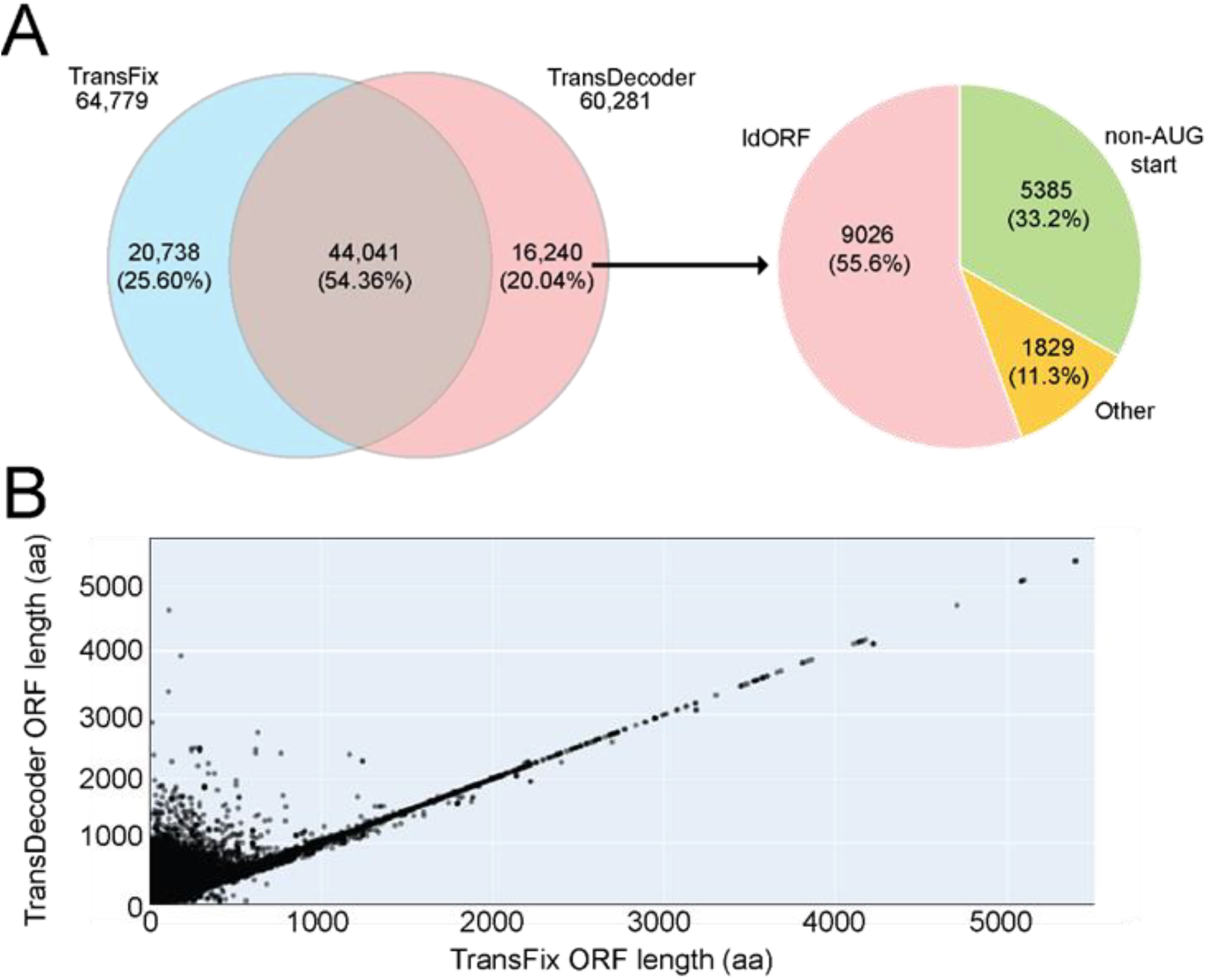
Comparison of ORF lengths from TransFix and TransDecoder for individual transcripts. **A)** Venn diagram of number of ORFs from TranSuite and TransFix – overlap represents ORFs with exact matches of co-ordinates. The 16,240 ORFs unique to TransDecoder are divided into ldORFs, ORFs with non-AUG start codons and other (predicted fusion ORFs, translations of opposite strand). **B)** Plot of ORF lengths generated for the same transcripts by TransDecoder (y-axis) and TransFix (x-axis).

The fundamental difference between ORFs generated by TransFix and TransDecoder is illustrated for *VERNALISATION2* (*VRN2*) (Figure 8). In TAIR10, *VRN2* had two isoforms: the .1 isoform which codes for the full-length protein and the .2 isoform which has an intron retention event of intron 2 (I2R) which altered the reading frame. The annotated ORF of the .2 isoform in TAIR begins at an AUG in exon 4. TransFix “fixed” the translation start site at that used to translate the .1 isoform resulting in a PTC and a short ORF for the .2 transcript. AtRTD2 contains seven transcript isoforms for this gene, of which P1 (same as TAIR.1) and s1 generate full-length protein variants differing by an alternative 3’ splice site in intron 11 that adds 6 nt (2 amino acids) (Figure 8). The remaining five transcript isoforms (c1, ID7, ID2, ID4 and .2) contained PTCs, are likely NMD substrates and were less abundant than the protein coding variants. TransFix translated all of the transcripts from the same translation start site generating three different PTC positions among the transcripts. In contrast, TransDecoder selected ldORFs downstream of the PTCs such that all translations ended at the same (authentic) stop codon but translation started at four different sites downstream of the authentic AUG (Figure 8). All seven transcript isoforms contained the translation start site used to translate the full-length protein variants (P1 and s1) and the fundamental principle of TransFix is that the ribosome will use the authentic translation start site instead of a downstream site that happens to produce an ORF longer than 100 amino acids (Brown et al., 2015).

**Figure 8.**
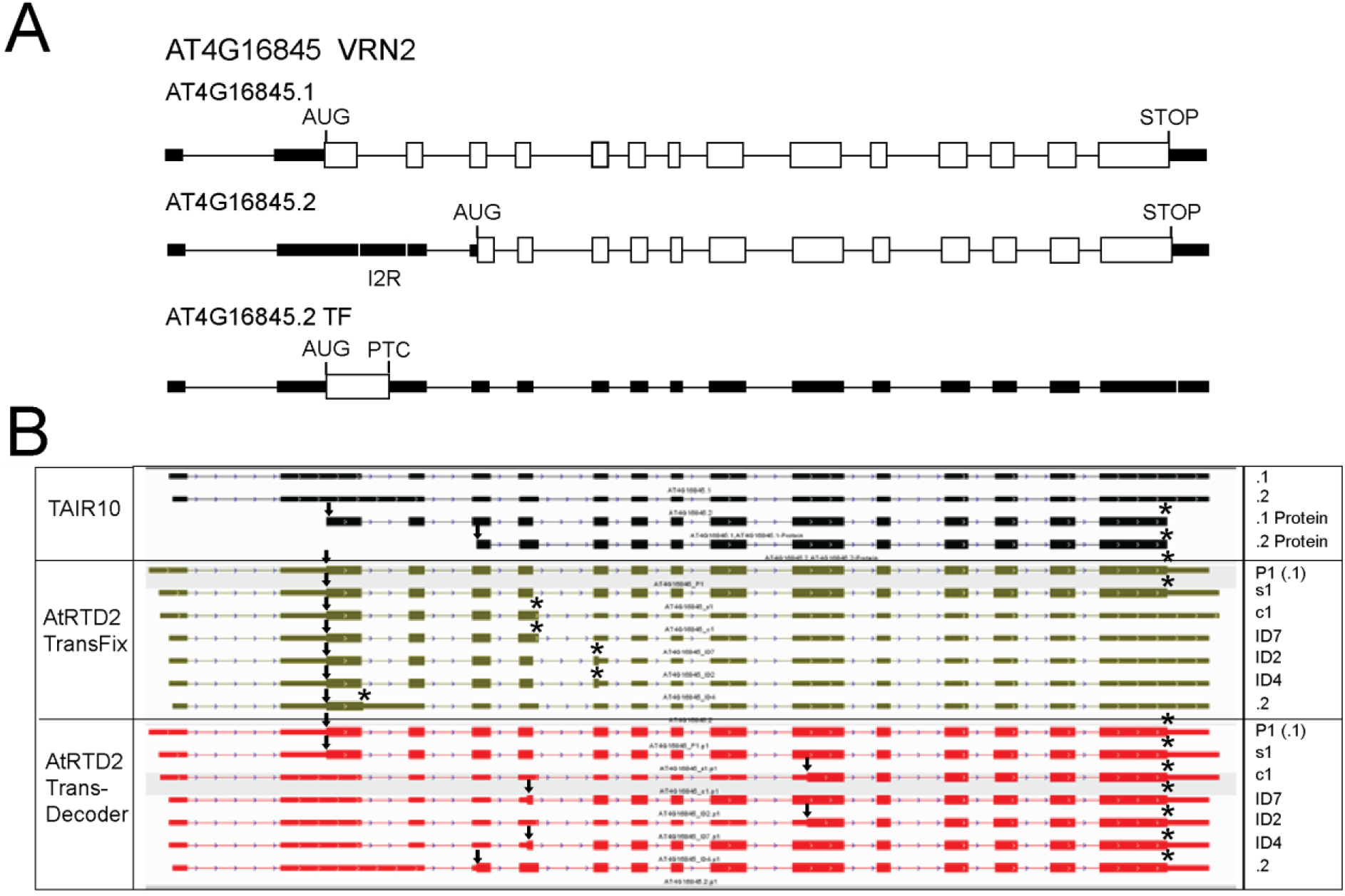
Comparison of TransFix and TransDecoder for Arabidopsis VRN2 gene isoforms. **A)** Transcript models (.1 and .2) from TAIR10 with annotated translations (.1 Protein and .2 Protein) and transcript model/translation of AT4G16845.2 from TransFix (TF). **B)** Screenshot from Integrative Genomics Viewer (IGV) of transcript models and ORFs from TAIR10 transcripts (top, black), and AtRTD2 transcript isoforms from TransFix (middle, green) and TransDecoder (bottom, red). In A) White boxes – coding exons; black boxes – UTRs. In B) arrows – translation start sites, asterisks – translation end sites; in IGV – thick exons – coding; narrow exons - UTRS.

We next investigated examples where the ORF length from TransDecoder was either longer or shorter than that from TranSuite (Figure 7B). We identified mis-calling of ldORFs and ORFs with N-terminal extensions from the other reading frames fused to C-terminal fragments (Supplementary Figures 1 and 2). In addition, some predicted ORFs from TransDecoder had non-AUG start codons (Supplementary Figures 2-4) and, in some cases, TransDecoder wrongly translated the opposite strand (Supplementary Figure 5). We therefore investigated the nature of the 16,240 (26.94%) ORFs unique to TransDecoder (Figure 7A). By comparing predicted ldORFs from TranSuite with ORFs from TransDecoder, TransDecoder had selected ldORFs in over 9,000 transcripts (Figure 7A; Figure 8). We also investigated the frequency of non-AUG start codons and found that a further 5,385 transcripts had non-AUG start codons (Figure 7A; Supplementary Figure 2). The non-AUG start codons represented all possible codons with a predominance of AAA lysine codons (Supplementary Figure 3). To understand the high frequency of non-AUG start sites, we examined examples of ORFs with non-AUG start codons and found that they consisted of a C-terminal fragment of the protein coded by the gene fused to an N-terminal extension derived from one of the other two reading frames that extended to the 5’-end of the transcript giving the non-AUG start codons (Supplementary Figures 2 and 4; Figure 3B.4). The N-terminal extensions to the end of the transcript varied in length in different genes and occurred irrespective of whether the gene contained introns, was alternatively spliced or whether transcripts contained PTCs. For example, some single exon genes (not shown) and multi-exon genes with only a single transcript showed extensions to the 5’-end of the transcript (Supplementary Figure 4A and B). These ORF arrangements appear to be a consequence of the TransDecoder program which prioritises ORFs in the first reading frame (the frame of the identified ORF) and ORF length and reflects the AS-induced frame switch causing (by chance) one of the other two frames to be in frame with a C-terminal ORF downstream of the PTC. Other genes also had N-terminal extensions which started at an AUG within the transcript (Figure 3B.2 and .3; Supplementary Figure 1C). From examples of genes which had some transcripts with an internal AUG and other transcripts where the ORF extended to the 5’-end of the transcript, it was clear that different outcomes depended on the presence or absence of an in frame stop codon between the identified ldORF and the 5’-end of the transcript, respectively (Supplementary Figures 2 and 4C and D). TransDecoder reports ORFs extending to the 5’-end as incomplete and missing a start codon (see TransDecoder website). The remaining ca. 2k ORFs unique to TransDecoder (Figure 7A) were fusion proteins of C-terminal fragments of the protein coded by the gene and unrelated N-terminal extensions with an AUG upstream of the PTC (see Figure 3B.3; Figure 8) or were translations from the opposite strand (Supplementary Figure 5). Thus, TransDecoder generates thousands of incorrect ORFs and for AtRTD2 around 27% of its transcripts would result in false ORFs.

## DISCUSSION

Robust translation and transcript characterisation programs are needed to exploit the mass of data in new and existing transcriptomes of different species, strains, cultivars, ecotypes and individuals. With the diversity of transcript isoforms generated by alternative splicing, it is also necessary to distinguish those that code for protein variants from unproductive isoforms which contain PTCs and are likely NMD targets. TranSuite is a complete pipeline of modules which generates true translations from transcripts. The basic tenets of TranSuite are three-fold. Firstly, for any gene the authentic translation start site is recognised by ribosomes to translate the primary protein of the gene. Secondly, any transcript from that gene which contains the authentic translation start site will be translated from that position because ribosomes have no mechanism to ignore the authentic translation start site that they normally use, in favour of a downstream AUG which might give a protein of >100 amino acids. Thirdly, translating from the authentic translation start site identifies transcripts with PTCs and on the basis of the position of the first PTC, a range of putative NMD features can be defined. In contrast, translation programs which select ORFs longer than a threshold (usually 100 amino acids) correctly identify ORFs for many genes and transcripts. However, for AS isoforms with frame shifts and PTCs, long downstream ORFs are often erroneously identified leading to frequent mis-annotation of ORFs in databases. TranSuite is also able to deal with more complex gene regulation arrangements such as genes with different transcription start sites and genes where AS has removed the primary AUG. The key advantage of TranSuite is that new or existing transcriptomes can be analysed for accurate translations and transcript features in a single pipeline.

The direct comparison between TranSuite and TransDecoder using Arabidopsis AtRTD2 (Zhang et al., 2017) showed that around 75% of ORFs were identical. The major difference was the identification of shorter ORFs ending in a PTC by TranSuite rather than ldORFs by TransDecoder. This difference was expected from the different methods of translation. However, TransDecoder also produced over >5k ORFs beginning with non-AUG codons due to artificial extensions of ORFs to the 5’-end of the transcript. Although translation can occur from non-AUG codons (Ivanov et al., 2008; Kearse and Willusz, 2017; Merchante et al., 2017) and such codons have been described in Arabidopsis (Simpson et al., 2010; Reichmann et al., 1999), the number of transcripts involved and the range of different non-AUG start codons from TransDecoder highlights the consequence of extending an ORF because it is in frame rather than translating from the authentic start site as in TranSuite. The artefactual non-AUG start codons above raise the question of real non-AUG start codons. Given the rarity of *bona fide* non-AUG start codons, it would not be possible to relax the requirement for an AUG in TranSuite to capture rare non-AUG start codons without generating vast numbers of false ORFs. Similarly, TranSuite is unable identify cases of reinitiation of translation, leaky ribosome scanning or use of IRES. For those genes with demonstrated non-AUG codons or other translation mechanisms, users can choose to manually alter their protein sequences.

Many AS transcripts from protein-coding genes are unproductive and prediction of whether they are targets of NMD is another goal of transcript characterization. TranSuite detects and annotates PTCs allowing downstream analyses of NMD signals including downstream splice junctions and long 3’UTRs. In the analysis of AtRTD2, three-quarters (76%) of the unproductive transcripts had the classic combination of NMD signals (PTC, DsSJ, long 3’UTRs) and a further 16% had at least two of these signals and/or an overlapping uORF suggesting that they are likely targets of NMD. Defining translation start and stop sites also helps to define 5’ and 3’UTR regions of genes/transcripts correctly allowing identification of uORFs. In Arabidopsis, we found previously that transcripts with overlapping uORFs correlated with NMD (Kalyna et al., 2012). In frame uORFs of 30-50 amino acids upstream of the AUG translation start triggered NMD (Nyiko et al., 2009) but also a much shorter uORF of only 13 amino acids induced NMD and inhibited translation of the main ORF (Saul et al., 2009). Thus, the rules governing the characteristics of uORFs that trigger NMD are not well defined. TranSuite identifies overlapping uORFs as potential NMD signals (Kalyna et al., 2012) and although it is unlikely that the majority of uORFs in 5’UTRs will trigger NMD, TranSuite flags all uORFs which code for peptides of >10 amino acids and provides information on whether the uORFs are in frame or out of frame with the main protein-coding ORF or are overlapping.

Alternative splicing is important in regulation of expression and AS in the 5’ and 3’ UTR can have important regulatory roles in controlling mRNA turnover or export. While characterising the AtRTD2 transcriptome from Arabidopsis, we found that AS was prevalent in UTRs with over 14k transcripts (26.1% of protein-coding transcripts) having AS in the 5’ and/or 3’UTR. AS in the 5’UTR can affect the position of the translation start site, size/position/frame of uORFs and other post-transcriptional and translational regulatory elements (Kalyna et al., 2012). Introns in the 3’UTR can alter transcript stability by, for example, triggering NMD because they produce a splice junction downstream of the authentic stop codon such that splicing or non-spicing can affect whether a transcript is degraded by NMD or not. In addition, in plants, transcripts that contain a retained intron avoid NMD because they are detained in the nucleus (Marquez et al., 2012; Kalyna et al., 2012; Göhring et al., 2014). Gene regulation by detained introns controlling mRNA degradation, export and translation are now common in eukaryotic systems (Jacob and Smith, 2017). We have identified numerous Arabidopsis genes where AS changes the levels of transcripts with intron retention in the 5’ or 3’UTR in response to cold (James et al., 2012; Calixto et al., 2018). The consequences of retention or splicing of 5’ and 3’UTR introns on nuclear detention, mRNA stability and protein expression is an area that requires further attention.

To date, protein-coding capacity in gene sequences has been determined by identifying ORFs coding for proteins of >100 amino acids disregarding genes coding for smaller proteins. In Arabidopsis, many genes and gene families code for proteins of <100 amino acids (Lease and Walker, 2006). For example, the *CLAVATA3 (CLV3)/ENHANCER OF SHOOT REGENERATION (ESR)/*(*CLE*) gene family contains 24 genes coding for proteins of 80-120 amino acids (Cock and McCormick, 2001; Sharma et al., 2003). The *LEAF CURLING RESPONSIVENESS* (*LCR*) multigene family in Arabidopsis contains 29 genes, the majority of which code for proteins of 75-105 amino acids but one gene codes for a protein of only 59 amino acids (Vanoosthuyse et al., 2001). Similarly, the *SELF-INCOMPATIBILITY RESPONSE* (*SCRL*) family has 28 genes in Arabidopsis with most coding for proteins of 70-109 amino acids but one gene codes for a protein of 63 amino acids (Vanoosthuyse et al., 2001). The *INFLORESCENCE DEFICIENT IN ABS*CISSION (*IDA*) protein is 77 amino acids (Butenko et al., 2003) and *DEVIL* (*DVL*) is only 51 amino acids (Wen et al., 2004). Other gene families such as the rapid alkalinisation factor-like (RALFL) (Olsen et al., 2002) and lipid transfer protein (Arondel et al., 2000) also code for short proteins. TranSuite allows the threshold to be defined and we analysed AtRTD2 using either >100 or >70 amino acids as the threshold for calling a protein-coding gene. The main impact of the 70 amino acid limit was to increase the number of protein-coding genes and decrease the number of non-protein coding genes by around 2,000 genes and 2,800 transcripts. By determining translation start sites and cognate ORFs, TranSuite delivers more biologically meaningful translations, overcomes the issue of mis-annotated ORFs in databases and provides thorough transcript characterisation. The program is flexible, easy to use and will give a step change in quality of information for existing and newly emerging transcriptomes.

## DATA AVAILABILITY

TranSuite was developed with Python 3.6 and the Biopython package (version 1.72). It runs on Windows and Linux machines with any average system specifications (8GB Memory RAM and processor Intel Core i5, or equivalent). TranSuite is available in on our GitHub repository: https://github.com/anonconda.

