## Supplementary Figures for "TranSuite: a software suite for accurate translation and characterization of transcripts"

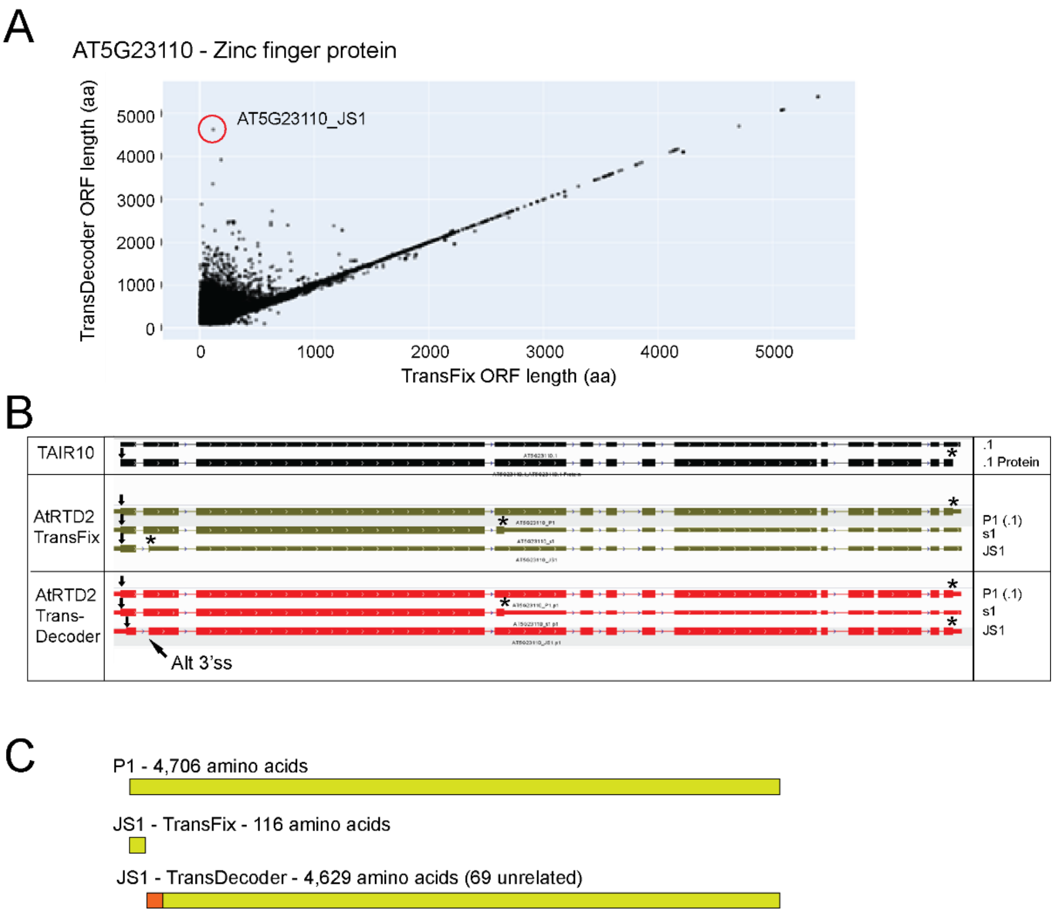


**Supplementary Figure 1. TransDecoder - Selection of in frame AUG generates false long ORF. A)** Plot of ORF lengths from each transcript using TransFix and TransDecoder. The AT5G23110_JS1 transcript has one of the largest differences in ORF length between TransFix and TransDecoder. **B)** Screenshot from IGB of transcript and translation models of AT5G23110 from TAIR10 (black) and AtRTD2 with TransFix (green) and TransDecoder (red). The JS1 transcript has an alternative 3' splice site (Alt 3'ss) in exon 2 which removes 110 nt, changes reading frame and introduces a PTC in exon 2 (TransFix JS1). **C)** Schematic diagram of predicted proteins from P1 and _JS1 transcripts. P1 codes for the full-length protein of 4,706 amino acids in both Transfix and TransDecoder. For the _JS1 transcript: in TransFix, the ORF is only 116 amino acids; in TransDecoder, the ORF is 4,629 amino acids consisting 69 amino acids from another frame fused to the 4,560 C-terminal amino acids of the normal protein. Arrows – translation start sites; asterisks – translation end sites; in IGV – thick boxes – coding regions; narrow boxes – UTRs; in C) green boxes – protein sequence of AT5G23110; orange box – unrelated protein from another reading frame.

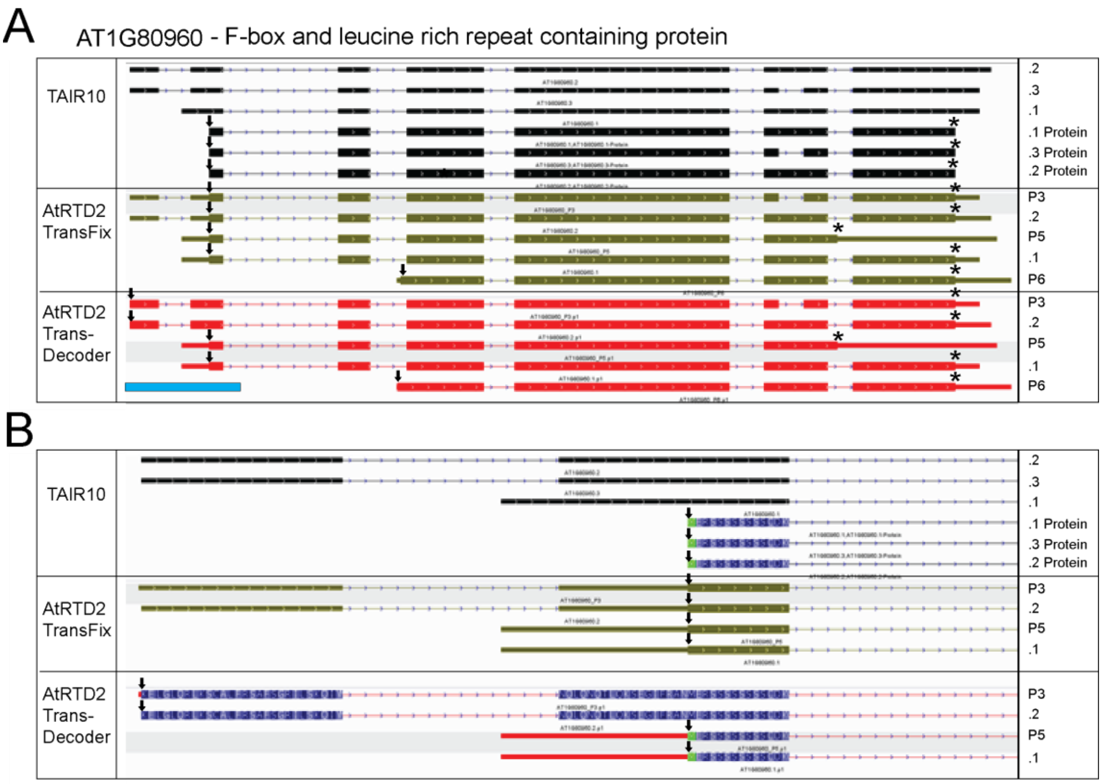


C

AT1G80960_P5 and .1

TransFix MERSSSSSSSSCDK

TransDecoder ENLVLSS*NQLQNQTLQKSEGIFRAN MERSSSSSSSSCDK

in frame stop codon

AT1G80960_P3 and .2

TransFix MERSSSSSSSSCDK

TransDecoder

KELGLQRLKSCALFRSAFSGRILSKQIY//NQLQNQTLQKSEGIFRAN MERSSSSSSSSCDK

no in frame stop codon

**Supplementary Figure 2. TransDecoder - Selection of non-AUG translation start codons generates false long ORF. A)** Screenshot from IGV of transcript and translation models of AT1G80960 from TAIR10 (black) and AtRTD2 with TransFix (green) and TransDecoder (red). TAIR10 has three AS isoforms with the same translation start and end points. AtRTD2 has five AS isoforms: P3 and .2 have an intron in the 5'UTR, P5 and .1 have an alternative transcription start site in intron 1 and P6 an alternative transcription start site in intron 3. In TransFix the four AS isoforms (P3, .2, P5 and .1) use the authentic translation start AUG and the .2 transcript has a different stop codon in the retained last intron. In TransDecoder, P5 and .1 have the same AUG start site as in TAIR10 and TransFix but the P3 and .2 ORFs start at a lysine residue (AAA) at the 5-end of the transcript adding 46 amino acids to generate an incorrect N-terminally extended ORF. **B)** Enlargement of 5' ends of transcripts and N-terminal ends of ORFs covering region marked by blue box in A) showing N-terminal ORF sequence differences between P3/.2 and P5/.1. **C)** Translations of 5'-ends of transcripts. Both TransFix and TransDecoder produce the correct translation for the P5 and .1 transcripts (red); the presence of an in frame stop codon is noted. TransFix produces the correct translation for P3 and .2 (red) but TransDecoder produces ORFs with an N-terminal extension (boxed) to the normal protein sequence (red); the extension goes to the 5'-end of the transcript with no in frame stop codon (// - intron position). Arrows – translation start sites; asterisks – translation end sites; in IGV – thick boxes – coding regions; narrow boxes - UTRs.

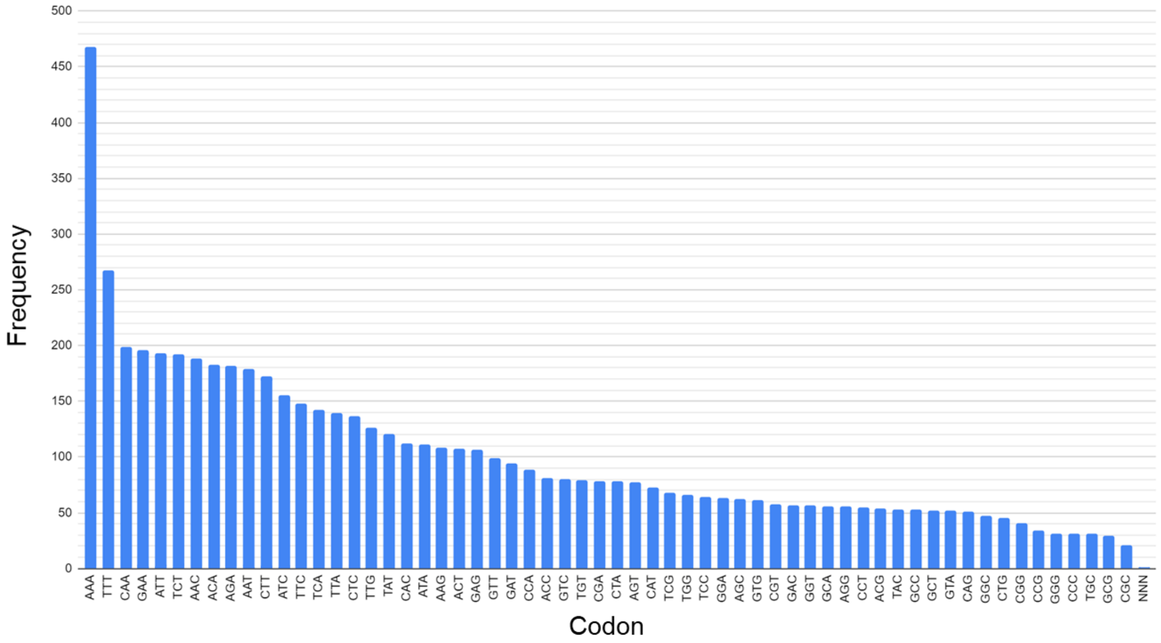


**Supplementary Figure 3. TransDecoder - Frequency of occurrence of non-AUG start codons among TransDecoder ORFs.**

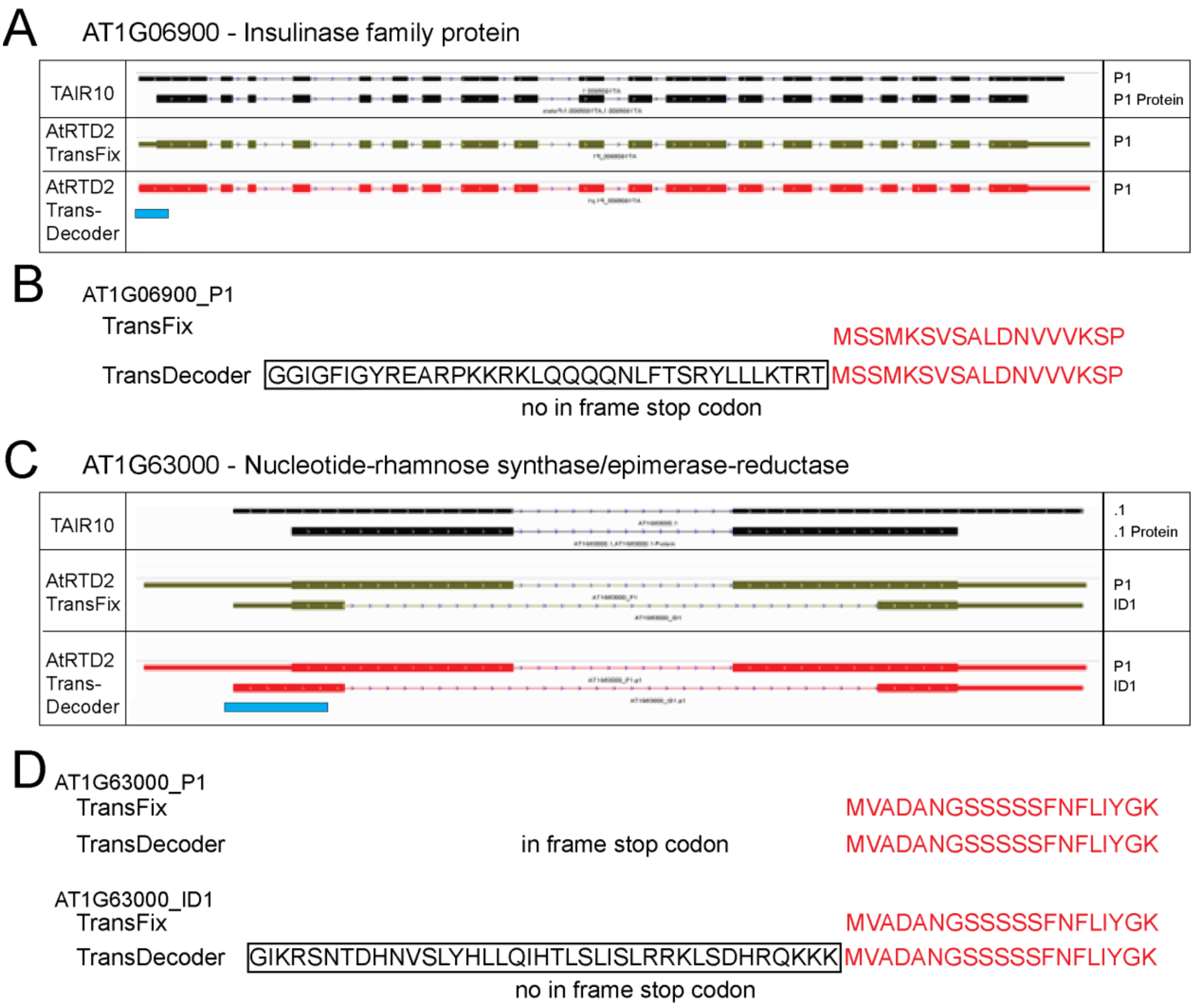


**Supplementary Figure 4. TransDecoder – False ORFs with non-AUG start codons. A)** Screenshot from IGV of transcript and translation models of AT1G06900 from TAIR10 (black) and AtRTD2 with TransFix (green) and TransDecoder (red). AT1G06900 is a multi-exon gene with a single transcript (P1). **B)** Protein sequences of ORFs covering the area indicated by the blue line in A). TransFix produces the correct ORF (red) but TransDecoder produces an ORF with an N-terminal extension (boxed) to the normal protein sequence (red); the extension goes to the 5'-end of the transcript with no in frame stop codon. **C)** AT1G63000 has two transcripts in AtRTD2 (P1 and _ID1) where ID1 has a shorter 5' UTR and larger intron. **D)** Both TransFix and TransDecoder produce the correct translation for the P1 transcript (red); the presence of an in frame stop codon is noted. TransFix produces the correct translation for ID1 (red) but TransDecoder produces an ORF with an N-terminal extension (boxed) to the normal protein sequence (red); the extension goes to the 5'-end of the transcript with no in frame stop codon. In IGV – thick boxes – coding regions; narrow boxes - UTRs.

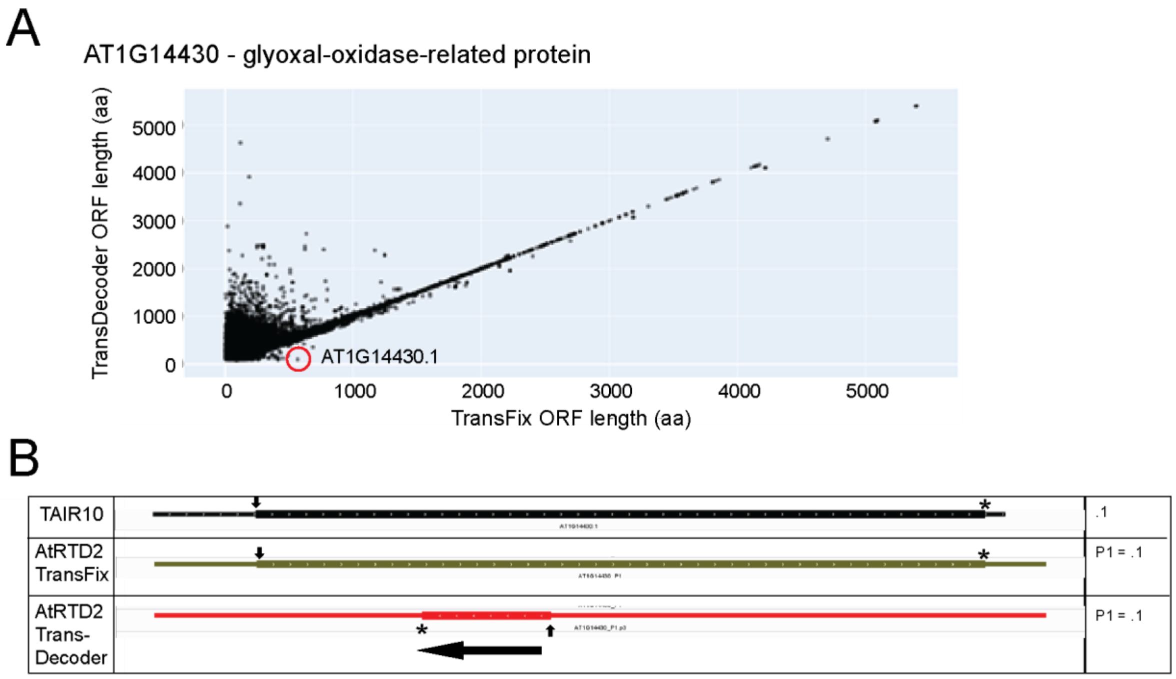


**Supplementary Figure 5. TransDecoder - Selection of ORF from opposite strand.** A) Plot of ORF lengths from each transcript using TransFix and TransDecoder. The AT1G14430.1 transcript has a much smaller ORF in TransDecoder than TransFix. B) Screenshot from IGV of transcript/translation model of AT1G14430.1 from TAIR10 (black) and AtRTD2 with TransFix (green) and TransDecoder (red). Arrows – translation start sites; asterisks – translation end sites; in IGV – thick boxes – coding regions; narrow boxes – UTRs; black arrow – direction of translation in TransDecoder.
